# Metabolic heterogeneity coupled with a resistance gene generates antibiotic heteroresistance

**DOI:** 10.1101/2025.03.11.641639

**Authors:** Jacob E. Choby, David A. Hufnagel, Muqing Ma, Tugba Ozturk, Victor I. Band, Minsu Kim, David S. Weiss

## Abstract

Heteroresistance is a form of antibiotic resistance in which a phenotypically unstable subpopulation of resistant cells exists within a majority population of susceptible cells. The impact of the extracellular environment on mechanisms of heteroresistance is unclear. Studying fosfomycin heteroresistance in an *Enterobacter cloacae* complex isolate, we observed that glucose availability greatly increased the frequency of the resistant subpopulation. Glucose downregulated the glycerol and fosfomycin importer GlpT, whose expression was heterogenous at the single cell level. This heterogeneous expression of GlpT, in combination with the expression of the fosfomycin resistance gene *fosA*, which acted as a resistance enhancer, led to the generation of heteroresistance. Correspondingly, the frequency of the fosfomycin resistant subpopulation was increased in murine models of hyperglycemia/diabetes. These data demonstrate how metabolic heterogeneity and carbon source availability can impact antibiotic resistance phenotypes in the infection environment.

## INTRODUCTION

Bacterial infections are a major cause of global morbidity and mortality. The rise and spread of antibiotic resistance among pathogens is a growing challenge to preventing and treating bacterial infections^1^. Bacteria may develop resistance through genetically encoded resistance determinants which include^2^: intrinsic resistance owing to the identity of the bacterium (i.e. the outer membrane of Gram-negative bacteria is resistant to many antibiotics), mutations to the gene encoding the antibiotic target, efflux pumps, or horizontally transmitted genes which encode enzymes that inactivate the antibiotic or modify the antibiotic target (i.e. β-lactamases). Conventionally, a clonal population from a single bacterial isolate which does not genetically encode resistance is entirely killed by the antibiotic and the isolate is considered susceptible. Conversely, a resistant isolate typically encodes some resistance determinant, and the entire clonal population survives antibiotic exposure. Clinical microbiology uses antimicrobial susceptibility testing to define isolates as susceptible or resistant to various antibiotics to advise antibiotic therapy.

Heteroresistance is a unique form of antibiotic resistance which defies conventional classification. A form of phenotypic heterogeneity, heteroresistant (HR) isolates have a minor subpopulation of resistant cells in a large majority susceptible population^3^. This resistant subpopulation is often challenging to detect by traditional antimicrobial susceptibility testing. The resistant cells grow in the presence of antibiotic, distinguishing them from a persister population, and antibiotic exposure will transiently enrich for the resistant subpopulation. When the antibiotic is removed, the phenotypically resistant population will return to baseline minority levels. How the resistant subpopulation behaves during human antibiotic therapy remains a major focus, as heteroresistance has been proposed as a cause of unexpected antibiotic treatment failure^4^.

Heteroresistance serves as a model of bacterial phenotypic heterogeneity; understanding the mechanisms that generate HR will inform bacterial cell biology as well as antibiotic therapy in the clinic. One mechanism by which bacterial populations exhibit HR is amplification of antibiotic resistance genes^5^. Gene amplification is a dynamic and reversible process by which a region of the genome is present in multiple copies, and copy number can be heterogenous in the population. Thus, amplification of an antibiotic resistance gene in a minor subpopulation confers greater resistance to the antibiotic^6,7^. Gene amplification has been described as the cause of HR in a variety of species to various antibiotics, and appears to be the most frequently identified mechanism of HR in Gram-negative clinical isolates^3,6–8^. On the other hand, gene amplification is not the sole cause of HR. For example, amplification-independent mechanisms have been described for colistin heteroresistance^9–12^. In this study, we sought to understand the mechanism of fosfomycin HR.

Fosfomycin was discovered over fifty years ago and is now increasingly used as oral therapy for urinary tract infections^13^ and there is renewed interest in its utility as an intravenous treatment for multi-drug-resistant infections. Fosfomycin inhibits peptidoglycan synthesis by covalently binding and inactivating the UDP *N*-acetylglucosamine enolpyruval transferase, MurA^14^. Fosfomycin HR is widespread in carbapenem-resistant Enterobacterales (CRE); we previously screened CRE clinical isolates through the Georgia Emerging Infections Program’s Multi-site Gram-negative Surveillance Initiative and observed that fosfomycin HR was detected at a greater rate (72% of isolates) than HR to any other antibiotic tested^15^.

Fosfomycin resistance primarily occurs through mutations to the fosfomycin target MurA, fosfomycin importers, or acquisition of a fosfomycin resistance determinant^16^. Fosfomycin enters the cell through glycerol-3-phosphate (GlpT) or hexose phosphate (UhpT) transporters, therefore mutations that inactivate this function can confer resistance by blocking fosfomycin entry^17–19^. Acquisition of the glutathione-S-transferase FosA also confers resistance; FosA conjugates glutathione to fosfomycin to inactivate the drug^20^. Various chromosomal and plasmid-borne *fosA* variants exist and are common among *Enterobacter* and *Klebsiella* clinical isolates but do not appear to have extensively spread into *Escherichia coli*^21^.

While much is understood about routes to fosfomycin resistance, what generates population-wide heterogeneity in fosfomycin resistance was not understood. In this study, we investigated fosfomycin HR in a model CRE isolate, *Enterobacter* strain Mu208. We found that the minority resistant subpopulation is supported by FosA and synthesis of the FosA cofactor, glutathione, confirming their role in fosfomycin resistance. Surprisingly, we found that the population heterogeneity was not the product of gene amplification, but rather heterogeneity in the activity of the *glpT* promoter, encoding the fosfomycin importer. Cells with low *glpT* expression constitute the resistant subpopulation. A transcriptional repressor GlpR and transcriptional activator CRP both play important roles in regulating *glpT* expression and responding to environmental carbon sources. Notably, glucose import repressed *glpT* expression via CRP and thereby increased population-wide fosfomycin resistance. This was recapitulated in murine models of hyperglycemia/diabetes; the frequency of the fosfomycin resistant subpopulation was enhanced in the presence of excess glucose during infection. Together, our data provide an example of amplification-independent heteroresistance, in which heterogeneity of an antibiotic importer results in heterogeneity of resistance. Because the metabolic environment regulates the importer expression, the context of infection may dictate the outcome of treatment of fosfomycin heteroresistant isolates.

## RESULTS

### *Enterobacter cloacae* complex isolate Mu208 is fosfomycin heteroresistant and causes antibiotic treatment failure in mice

We investigated the mechanism of fosfomycin heteroresistance (HR) in an *Enterobacter cloacae* complex strain isolated from a human infection, strain Mu208. To investigate HR, the population analysis profile (PAP; Supp. Fig. 1) is used to assess cells in the population that are able to form colonies on a range of antibiotic concentrations. The PAP assesses the population at concentrations above and below the clinical breakpoint. The breakpoint concentration differentiates resistant versus susceptible isolates in clinical testing for each antibiotic; the CLSI fosfomycin breakpoint is 256 µg/mL^22^. Mu208 exhibited fosfomycin HR, harboring a subpopulation resistant to 256 µg/mL fosfomycin and greater. The subpopulation constitutes approximately 1-10% of the total population at this concentration. This contrasts with the susceptible *Enterobacter* isolate Mu819 in which all the cells are uniformly killed at a concentration of fosfomycin below the clinical breakpoint (Figure 1a, Supp. Fig. 2a). In a time kill assay, there is a period of initial killing by fosfomycin but subsequent rapid growth of Mu208, and the growing population is comprised of fosfomycin resistant cells (Figure 1b). The resistant subpopulation which grows in the time-kill assay is also unstable: following enrichment of the resistant cells in broth containing fosfomycin, the resistant population returns to baseline frequency following growth in fosfomycin-free media (Figure 1c). Together, these data establish that strain Mu208 exhibits unstable fosfomycin heteroresistance.

Next, we tested whether the minority resistant subpopulation is sufficient to cause treatment failure in a murine model of peritonitis. As expected, fosfomycin treatment rescued mice from a lethal dose of the susceptible Mu819 isolate (Figure 1d). However, fosfomycin treatment did not rescue mice infected with Mu208, demonstrating that this HR isolate caused treatment failure (Figure 1e).

### The fosfomycin resistance enzyme FosA and synthesis of the FosA cofactor, glutathione, are required for the gene-amplification independent resistant subpopulation

We sought to understand the mechanistic basis for the phenotypic heterogeneity that results in fosfomycin heteroresistance. Amplification of antibiotic resistance genes has been found to generate an antibiotic-resistant subpopulation in a variety of species to various antibiotics^3,6–8^. We therefore tested whether the fosfomycin resistant subpopulation in Mu208 was a result of gene amplification, but our data did not support this hypothesis. First, the Δ*recA* strain, which is unable to perform RecA-dependent homologous recombination important for some gene amplification events, remains fosfomycin HR (Supp. Fig. 2b). Second, genome sequencing of resistant subpopulations did not identify areas of the genome with elevated gene copy number >2 except at the chromosome and plasmid origin (Supplemental Table 1). Copy number variation in the resistant subpopulation is a common tool to identify regions of gene amplification.

Therefore, we hypothesized that fosfomycin HR in Mu208 was amplification-independent and turned to a transposon screen to identify genetic factors that reduced the population resistance to fosfomycin. We screened ∼80,000 colonies following transposon mutagenesis for a loss in the ability to form a colony on agar containing fosfomycin (Supp. Fig. 3, Supplemental Table 2). Mu208 harbors a chromosomal *fosA2/fosA^EC^* family variant^21^ (hereafter referred to as *fosA)*, encoding a glutathione-S-transferase, which conjugates glutathione to the epoxide ring of fosfomycin, inactivating the drug. A validating hit, the fosfomycin resistance gene *fosA* was one gene that reduced fosfomycin resistance when disrupted. A cluster of transposon mutants with reduced fosfomycin resistance was associated with synthesis of the FosA cofactor glutathione (Figure 1f). We generated a Δ*fosA* strain which demonstrated that FosA was required for the highly resistant subpopulation (Figure 1g, Supp. Fig. 4a). Nevertheless, heterogeneity remained in the Δ*fosA* strain, with subpopulations that had a greater relative resistance than the majority population but did not survive to the breakpoint. This appears to be a common feature of Enterobacterales lacking *fosA*^23^, and also suggests FosA-independent heterogeneity. Interestingly, complementation *in trans* with a multi-copy plasmid resulted in homogenous resistance (100% of cells are phenotypically resistant) to even high concentrations of fosfomycin (Figure 1g). Similarly, inactivation of glutathione synthesis by generating the Δ*gshA* strain also reduced the frequency of the resistant subpopulation (Figure 1h, Supp. Fig. 4b). Exogenous glutathione increased the resistant subpopulation frequency (Supp. Fig. 4c), even in the absence of endogenous glutathione synthesis (Δ*gshA*), but the frequency remained much lower than wildtype (Supp. Fig. 4e). Exogenous addition of the glutathione precursor L-glutamate also increased the resistant subpopulation frequency (Supp. Fig. 4d). The effect of L-glutamate was dependent on endogenous glutathione synthesis (Supp. Fig. 4e). Therefore, the resistant subpopulation requires FosA and its cofactor glutathione.

We were intrigued by the contribution of endogenous glutathione synthesis to fosfomycin HR, and the availability of glutathione during infection. Glutathione is an important mammalian cofactor which can achieve millimolar intracellular and micromolar extracellular concentrations^24^. To test whether bacterial-derived glutathione is required even in the presence of mammalian-derived glutathione, we infected mice with a lethal dose of the Δ*gshA* glutathione synthase mutant, and found that this strain did not cause treatment failure (Figure 1i-j). Thus, synthesis of the FosA cofactor glutathione is critical to the fosfomycin resistant subpopulation and the ability of strain Mu208 to cause fosfomycin treatment failure *in vivo*.

**Figure 1.**
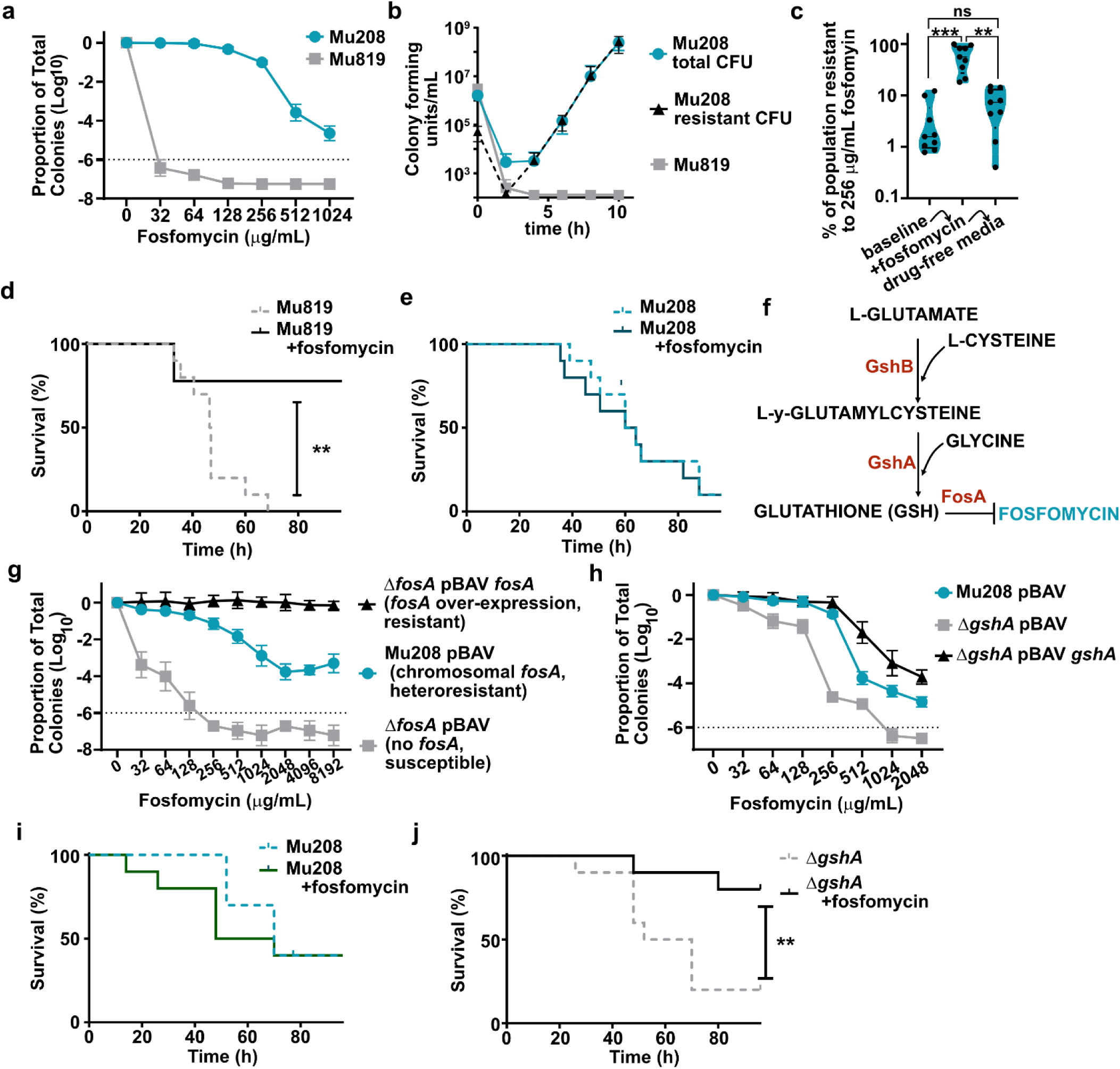
*Enterobacter cloacae* isolate Mu208 is fosfomycin heteroresistant and causes treatment failure in mice. (a) Population analysis profile (PAP) of Mu208 and *E. cloacae* Mu819 plated on Mueller-Hinton agar (MHA) containing fosfomycin; the proportion of surviving colonies is quantified relative to MHA containing 0 fosfomycin, from three independent experiments with 2 biological replicates each. (b) Time kill of Mu208 and Mu819: survival in media containing 512 μg/mL fosfomycin over time and quantification of surviving colony forming units on MHA (Mu208 total and Mu819) and MHA containing 512 μg/mL fosfomycin (resistant CFU) from two independent experiments with 5 biological replicates each. (c) Quantification of the fosfomycin resistant subpopulation of Mu208 after growth in media alone (baseline), subcultured into 256 μg/mL fosfomycin and grown for 20 h, and subcultured into fresh media without fosfomycin. At the end of each growth, an aliquot was diluted and plated onto MHA containing 256 μg/mL fosfomycin to quantify the resistant subpopulation, from three independent experiments with 3 biological replicates each. ** indicates p=0.0044 and *** indicates p=0.0004 by RM one-way ANOVA with Sidak’s multiple comparisons test, F (1.847, 14.78) = 20.57. (d-e) Survival of mice infected with (d) Mu819 or (e) Mu208, with and without fosfomycin treatment. Mice (n=10 per group, except n=9 for Mu819+fos) were infected intraperitoneally (IP) and fosfomycin was administered at 200 mg/kg, IP. ** indicates p=0.0028 by Log-rank (Mantel-Cox) test. (f) Schematic of glutathione biosynthesis and inactivation of fosfomycin, genes indicated in red were transposon hits. (g) PAP of Mu208, the Δ*fosA* deletion mutant, and its complement, from two independent experiments with 2 biological replicates each. (h) PAP of Mu208, a *gshA* deletion mutant, and its complement, from two independent experiments with 3 biological replicates each. (i-j) Survival of mice (n=10 per group) infected with (i) Mu208 or (j) Mu208 Δ*gshA*, with and without fosfomycin (fos) treatment. Mice were infected intraperitoneally (IP) and fosfomycin was administered at 200 mg/kg, IP. ** indicates p=0.0058 by Log-rank (Mantel-Cox) test.

### Expression of the fosfomycin importer GlpT is heterogenous and predicts survival in fosfomycin

Having established that Mu208 encodes the fosfomycin resistance determinant FosA, and can synthesize the FosA cofactor glutathione, it was unclear why the majority of the population was susceptible to the breakpoint concentration of fosfomycin. Thus, we turned to features that make Mu208 vulnerable to fosfomycin, reasoning that heterogeneity in those features could result in heteroresistance. Fosfomycin enters cells through the carbon importers GlpT (glycerol-phosphate) and/or UhpT (hexose phosphate). Mu208 encodes both, but *uhpT* has a frame-shift mutation rendering it nonfunctional (mutations in *uhpT* are prevalent among *Enterobacter* isolates^25,26^). We tested the role of each transporter in fosfomycin susceptibility. The Δ*glpT* strain was impervious to even very high concentrations of fosfomycin (Figure 2a, Supp. Fig. 5) suggesting it is the primary fosfomycin importer in this strain under conditions tested. On the other hand, the Δ*uhpT* strain demonstrated no change in fosfomycin HR, consistent with its frameshift annotation and confirming the primacy of GlpT as a fosfomycin importer in Mu208 (Supp. Fig. 6a). Compared to the Δ*glpT* strain, wildtype Mu208 had ∼5 logs of killing at high concentrations, suggesting *glpT* is expressed under these conditions. We complemented the *glpT* deletion *in cis* at the *attB* site under control of the native promoter (P*_glpT_*) or a constitutive promoter (P*_tet_*). When *glpT* was constitutively expressed, the majority of the population was killed at much lower fosfomycin concentrations compared to the strain complemented with the native promoter, suggesting transcriptional control of *glpT* has a large impact on the population’s relative fosfomycin resistance (Figure 2a, Supp. Fig. 5).

Thus, we hypothesized that population-level variation in GlpT activity could explain heterogeneity in fosfomycin resistance. We generated a strain encoding P*_glpT_mCherry* in the chromosome to monitor activation of the *glpT* promoter. Mu208 att::P*_glpT_mCherry* was imaged on agar pads and fluorescence was measured per cell. Fosfomycin was added and then we performed time-lapse microscopy to assess the fate of each cell during fosfomycin exposure. We found that there is population wide heterogeneity in P*_glpT_* activity prior to fosfomycin exposure, and that cells with lower P*_glpT_* are the cells that survive subsequent fosfomycin treatment (Figure 2b). To confirm this finding was unique to P*_glpT_* and rule out the possibility that the resistant cells were globally transcriptionally less active, we similarly measured P*_gshA_mCherry* and found that P*_gshA_* activity was similar between resistant and susceptible cells (Supp. Fig. 7).

**Figure 2.**
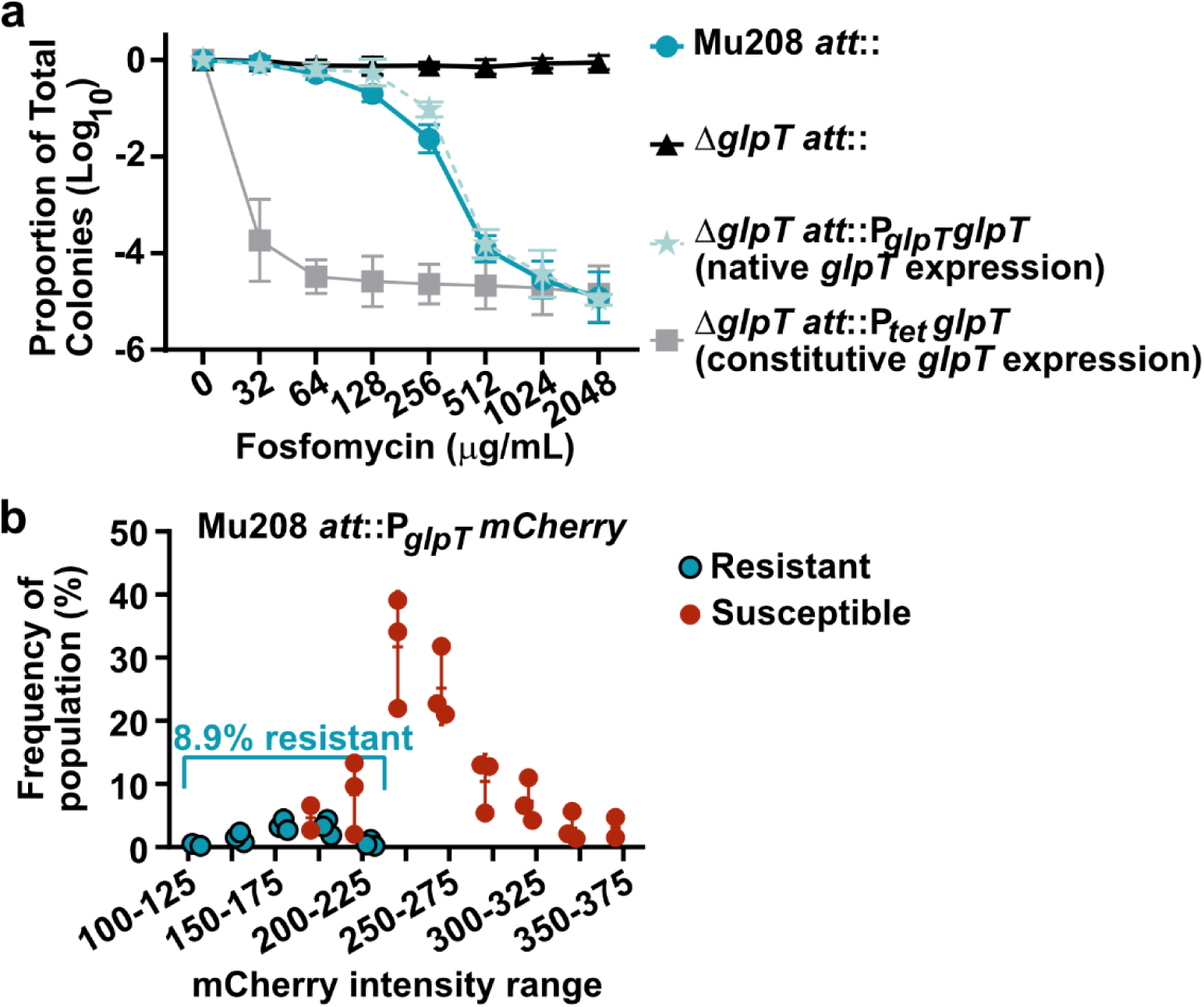
Expression of the fosfomycin and glycerol phosphate importer GlpT correlates with survival in fosfomycin. (a) PAP of Mu208, a *glpT* deletion mutant, and the *glpT* mutant complemented in the chromosome under control of its native promoter or a constitutive promoter, from two independent experiments with three biological replicates each. (b) P*glpTmCherry* expression intensity among cells that survived or were killed by 128 µg/mL fosfomycin, from three independent experiments; data were determined from 700-1000 cells for each experiment.

### *glpT* inactivation mutations occur at only a low frequency in the resistant subpopulation

Our data support a model that GlpT is the primary fosfomycin importer in Mu208 and expression of *glpT* is a critical determinant of relative fosfomycin resistance. Mutations which ablate the function of the fosfomycin importers GlpT could represent a route to fosfomycin resistance in our assays. Therefore, we evaluated the resistant subpopulation in Mu208 for changes in *glpT* function by sequencing or phenotypic testing. We sequenced colonies collected from agar containing fosfomycin and performed variant analysis. P*_glpT_glpT* mutations were undetected in 6/7 biological replicates (Supp. Fig. 6b). Phenotypically, fosfomycin resistant colonies were patched on M9 agar with glycerol phosphate as the sole carbon source, and colonies with null GlpT function were unable to grow. On 256 µg/mL fosfomycin, *glpT* null resistant colonies appeared at a rate of 1 per 100,000 total cells, much lower than the frequency of the total resistant subpopulation (Supp. Fig. 6c). While phenotypically null *glpT* mutants arise, they comprise only a small fraction of the resistant subpopulation at the breakpoint. However, when a strain only survives at a frequency of 1 per 100,000 total cells, or -5 logs because of genetic mutation or fosfomycin concentration, the majority of surviving colonies are phenotypically *glpT* null (Supp. Fig 6d).

### Transcriptional regulators GlpR and CRP control *glpT* expression and fosfomycin resistance

Returning to the importance of *glpT* regulation, we identified two regulator binding sites in P*_glpT_*: the transcriptional repressor GlpR^27^ and the carbon catabolite repression regulator CRP^28^ (cylic-AMP receptor protein) (Figure 3a). We generated deletion mutants of each regulator and assessed fosfomycin resistance by PAP. The Δ*glpR* strain was more susceptible to fosfomycin compared to wildtype Mu208, while the Δ*crp* strain was homogenously resistant, consistent with GlpR repressing P*_glpT_* and CRP activating P*_glpT_* (Figure 3b, Supp. Fig. 8a). These phenotypes were complemented *in trans* (Supp. Fig. 8b-c). We measured P*_glpT_mCherry* in the Δ*glpR* background and found that all cells in the population had high P*_glpT_* activity and were killed by fosfomycin (Figure 3c). Thus, GlpR repressed *glpT* expression to some extent in wildtype cells, affecting population wide fosfomycin resistance.

Next, we added exogenous cyclic adenosine monophosphate (cAMP) to Mu208 grown in minimal media and found that it reduced the frequency of the resistant subpopulation (Figure 3d) and increased *glpT* expression (Figure 3e). Additionally, inactivation of the cAMP-generating adenylate cyclase (*cyaA*) phenocopied the *crp* deletion (Supp. Fig. 8d). When P*_glpT_mCherry* was measured in the Δ*crp* strain, the entire population was resistant and had low P*_glpT_* activity (Figure 3f). We also tested whether the fosfomycin resistance which resulted from deletion of *crp* required the native *glpT* promoter: deletion of *crp* in a strain encoding the constitutively expressed *glpT* was as fosfomycin susceptible when *crp* was intact (Supp. Fig. 8e). Together, these data support the model that fosfomycin resistance is responsive to CRP activating expression of P*_glpT_.* We therefore tested whether the minor population that is fosfomycin resistant, which shows low P*_glpT_* activity (Figure 2b) might correlate with population-wide heterogeneity in CRP activity. We measured P*_crp_mCherry* and indeed observed heterogeneity amongst the population, with low P*_crp_* cells surviving fosfomycin treatment (Figure 3g). In sum, the carbon catabolite repression pathway exhibits heterogeneity, and there is a minor population of low CRP cells. These low CRP cells correlate with low *glpT*, as CRP is a transcriptional activator of P*_glpT_*. Low P*_glpT_* cells survive fosfomycin treatment because GlpT is the primary fosfomycin importer in strain Mu208.

**Figure 3.**
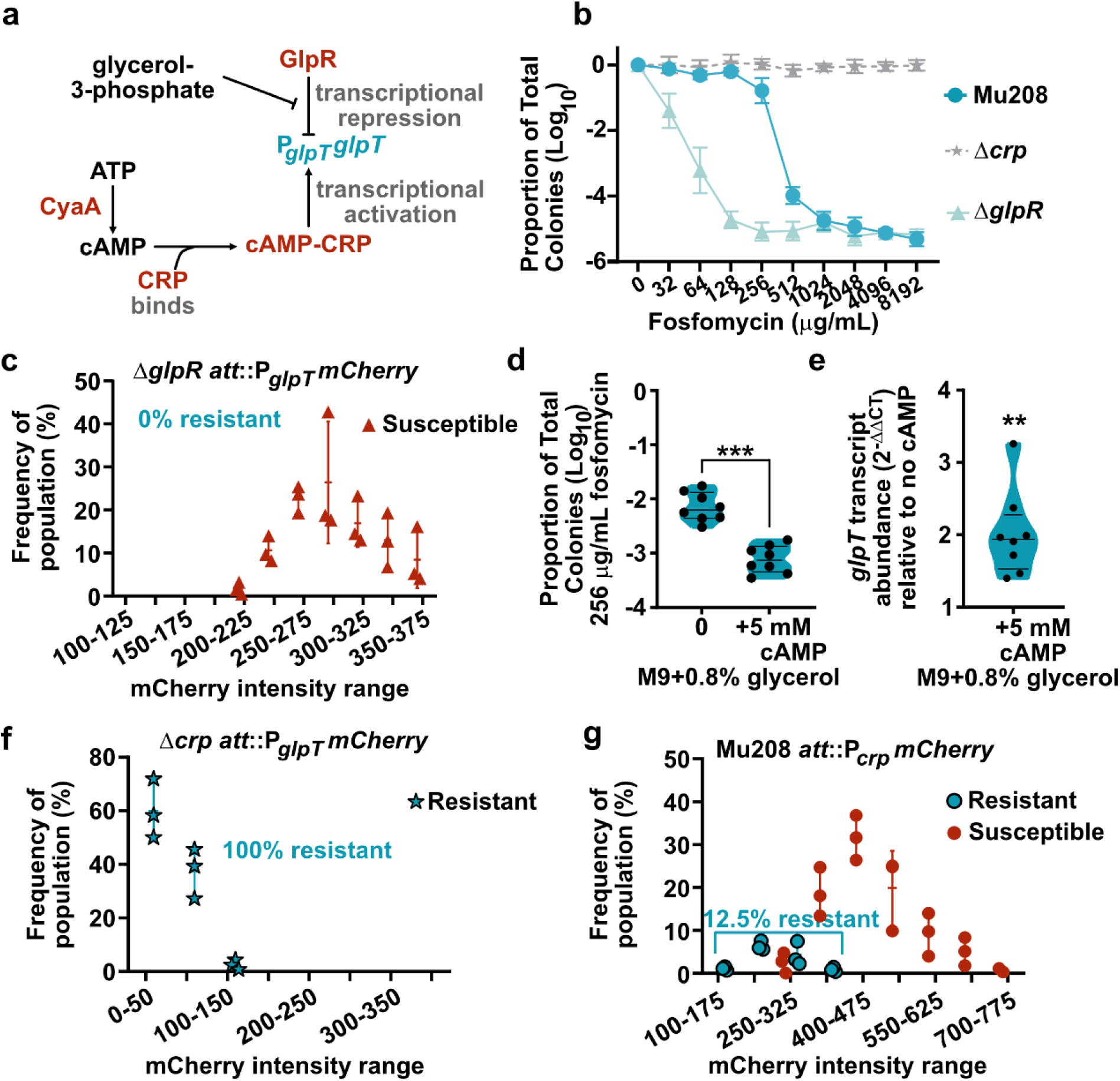
Transcriptional regulation of GlpT determines resistant subpopulation frequency. (a) Schematic of regulation of *glpT* and its promoter by 1] the GlpR repressor, binding of which is inhibited by the presence of glycerol-3-phosphate, and 2] CRP bound to cAMP, synthesized by CyaA. (b) PAP of Mu208 and mutants of cyclic AMP receptor protein (Δ*crp*), and glycerol phosphate repressor (Δ*glpR*), from two independent experiments with 3 biological replicates each. (c) *glpT* expression (P*glpTmCherry*) in the Δ*glpR* mutant and corresponding survival upon exposure to 128 µg/mL fosfomycin, from three independent experiments; data were determined from 700-1000 cells for each experiment. (d) Proportion of fosfomycin resistant colonies in the absence or presence of exogenous cyclic AMP from two independent experiments with 4 biological replicates each. *** indicates p= 0.0006 by two-tailed paired t-test, t=5.867, df=7. (e) Transcript abundance of *glpT* measured by qPCR in the absence or presence of exogenous cyclic AMP from C from two independent experiments with 4 biological replicates each, ** indicates p= 0.0019 by two-tailed paired t-test, t=4.816, df=7. (f) *glpT* expression (P*glpTmCherry*) in the Δ*crp* mutant and corresponding survival upon exposure to 128 µg/mL fosfomycin, from three independent experiments; data were determined from 700-1000 cells for each experiment. exposure. (g) *crp* expression (P*crpmCherry*) in Mu208 mutant and corresponding survival upon exposure to 128 µg/mL fosfomycin, from three independent experiments; data were determined from 700-1000 cells for each experiment.

### Glucose represses *glpT* expression via CRP and increases the frequency of the resistant subpopulation

The important role played by GlpR and CRP on *glpT* expression and fosfomycin resistance suggested that carbon source availability is a key regulator of fosfomycin HR. In support of this hypothesis, *ptsG* was identified in our transposon screen for loss of fosfomycin resistance (Supplemental Table 2). *ptsG* encodes the glucose-specific EIICB component of the sugar phosphotransferase (PTS) system; PtsG imports and phosphorylates glucose to generate intracellular glucose-6-phosphate. Therefore, we tested and found that exogenous glucose increased the frequency of the fosfomycin resistant subpopulation at the breakpoint concentration. Compared to wildtype, there was a ∼2 log reduction in the subpopulation frequency in Δ*ptsG*, and the Δ*ptsG* strain did not respond to exogenous glucose (Figure 4a). Import and phosphorylation of glucose by PtsG is known to reduce cAMP synthesis by CyaA, thereby reducing CRP activity^29^ and subsequent expression of *glpT*. We measured *glpT* transcript abundance by qPCR and found that excess glucose resulted in ∼100 fold reduction in relative *glpT* transcript abundance (Figure 4b).

Because of the key role of carbon source availability on fosfomycin resistance, we turned to M9 minimal media to control the carbon source present. When Mu208 was grown with glycerol as the sole carbon source, the frequency of the subpopulation resistant to 256 µg/mL was below 1%, but low concentrations of exogenous glucose resulted in a dose-dependent increase in frequency of the resistant population, which was not dependent on total carbon equivalents (Figure 4c). Additionally, the ability of glucose to enhance the resistant subpopulation appeared to be common in fosfomycin HR *Enterobacter* strains (Supp. Fig. 9). We hypothesized that the 2-log difference in resistant subpopulation frequency following growth in glycerol versus glucose as the primary carbon source was mediated largely by changes in *glpT* promoter activation. To test this, we measured the resistant population frequency of the strain encoding *glpT* under the P*_tet_* constitutive promoter and found that it was not responsive to glucose (Figure 4d), consistent with this hypothesis.

We performed PAP of Mu208, Δ*glpT*, Δ*gshA*, and Δ*fosA* strains following growth in M9 broth with glycerol or glucose. The entire population of wildtype Mu208 survived up to 64 µg/mL in M9 glycerol but up to 256 µg/mL in M9 glucose, with similar shifts for the Δ*gshA* and Δ*fosA* strains (Figure 4e-f, Supp. Fig. 10), consistent with glucose mediating a global effect on reduced fosfomycin entry by repressing *glpT* expression. We performed time-lapse microscopy with P*_glpT_mCherry* following growth in the same conditions and observed a shift in the peak fluorescence of the population from ∼250 in glycerol to <100 in glucose; the resistant subpopulation frequency shifted from 8% in glycerol to ∼95% in glucose (Figure 4g-h). These data are consistent with glucose enhancing fosfomycin resistance primarily through *glpT* repression and thereby limiting fosfomycin import.

**Figure 4.**
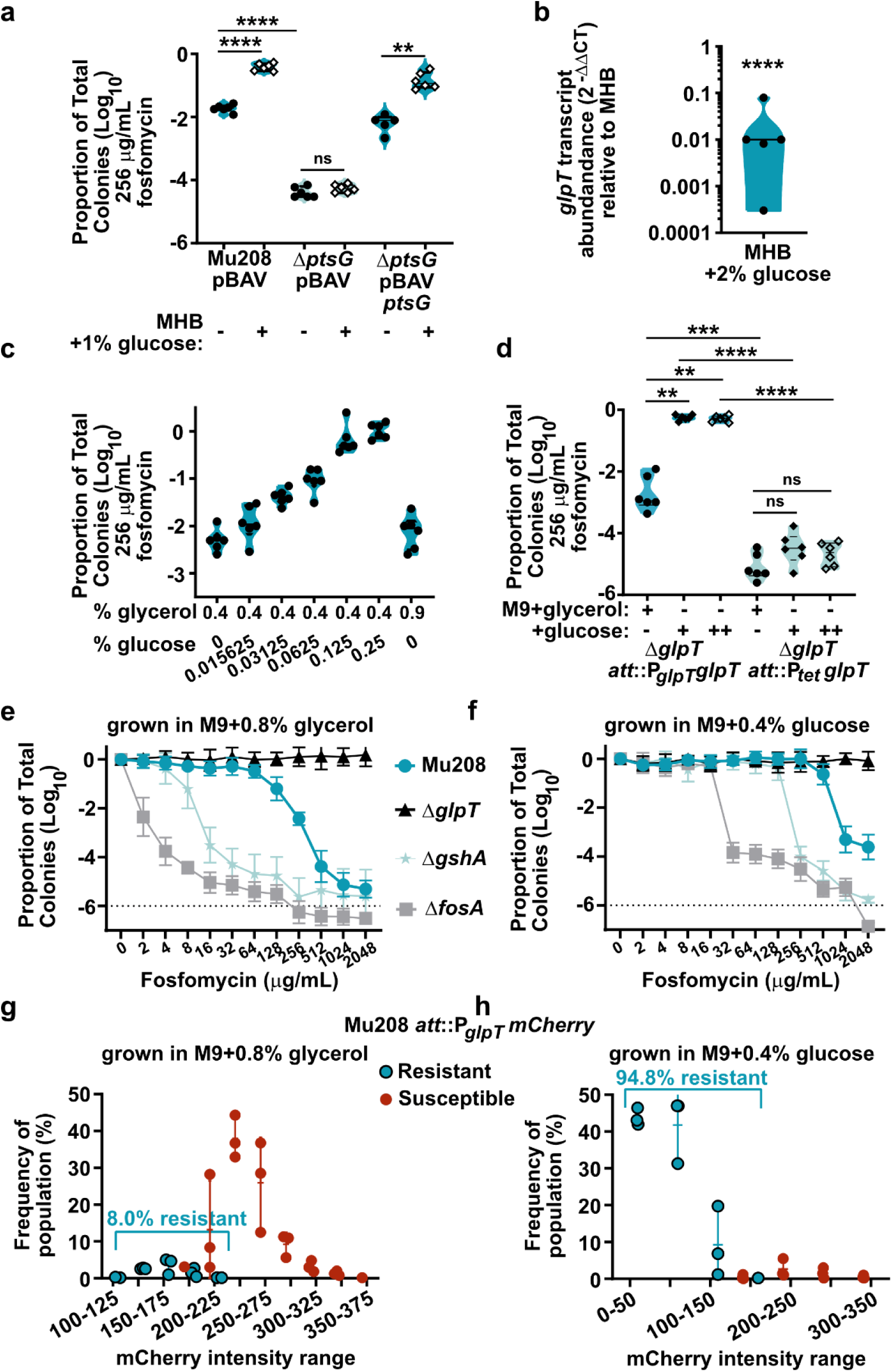
Glucose enhances the frequency of the resistant subpopulation by repressing *glpT*. (a) Proportion of colonies resistant to fosfomycin in Mu208, the Δ*ptsG* deletion mutant, and its complement, in Mueller Hinton broth with and without exogenous glucose, from two independent experiments with three biological replicates each. ns indicates p>0.05, ** indicates p=0.0027, and **** indicates p<0.0001 by mixed-effects analysis Geisser-Greenhouse correction and Šídák’s multiple comparisons test, F (2.471, 11.86) = 581.5 (b) *glpT* abundance measured by qPCR following growth in MHB+2% glucose relative to MHB alone, from two independent experiments with 5 biological replicates total. **** indicates p<0.0001 by two-tailed paired t-test, t=66.62, df=4. (c) Proportion of colonies resistant to fosfomycin in Mu208 grown in M9 media with glycerol and increasing concentration of glucose, from two independent experiments with 3 biological replicates each. (d) Proportion of colonies resistant to fosfomycin in Mu208 grown in M9 media with 0.8% glycerol, M9 with 0.6125% glucose (+) or M9 wth 2.5% glucose (++), from two independent experiments with 3 biological replicates each. ns indicates p>0.05, ** indicates p=0.0014 (+glucose), p=0.0011 (++glucose), *** indicates p=0.0008, and **** indicates p<0.0001 by RM one-way ANOVA with Šídák’s multiple comparisons test, F (2.402, 12.01) = 242.1. (e-f) Population analysis profiles from strains Mu208, Δ*glpT*, Δ*gshA*, and Δ*fosA*, (e) following growth in M9+glycerol as the sole carbon source or (f) following growth in M9+glucose as the sole carbon source, from two independent experiments with 3 biological replicates each. (g-h) *glpT* expression (P*glpTmCherry*) in Mu208 and corresponding survival upon exposure to 128 µg/mL fosfomycin in (g) M9+glycerol as the sole carbon source or (h) M9+glucose as the sole carbon source. From three independent experiments; data were determined from 700-1000 cells for each experiment.

### Excess glucose in murine models of diabetes increases the frequency of the resistant subpopulation

In our assays, changing the growth conditions resulted in substantial changes to the frequency of the resistant subpopulation. However, the mammalian infectious niche is not well-represented by growth conditions used for clinical susceptibility testing, or by M9 broth with a sole carbon source. Glucose levels in patients suffering from bacterial infection can vary based on co-morbidities, and diabetic patients are particularly vulnerable to bacterial infections^30^. Therefore, we sought to test whether excess glucose *in vivo* changed the frequency of the fosfomycin resistant subpopulation during infection. We used two models of hyperglycemia/diabetes: streptozotocin-treated mice to represent type I diabetes and the *ob/ob* obese mouse model of type II diabetes. These mice with their cognate controls were infected in the peritoneum with Mu208. Following 24 h of infection in the absence of fosfomycin treatment, a peritoneal wash and the spleen and liver (sites of dissemination) were collected. Samples were plated on agar containing 0 or the breakpoint concentration of fosfomycin to enumerate the frequency of the resistant subpopulation. In both models of excess mammalian glucose, the frequency of the population resistant to 256 µg/mL fosfomycin increased in at least two sites (Figure 5a-b, Supp. Fig. 11). These data suggest that during infection, Mu208 responds to host glucose and is more resistant to fosfomycin.

**Figure 5.**
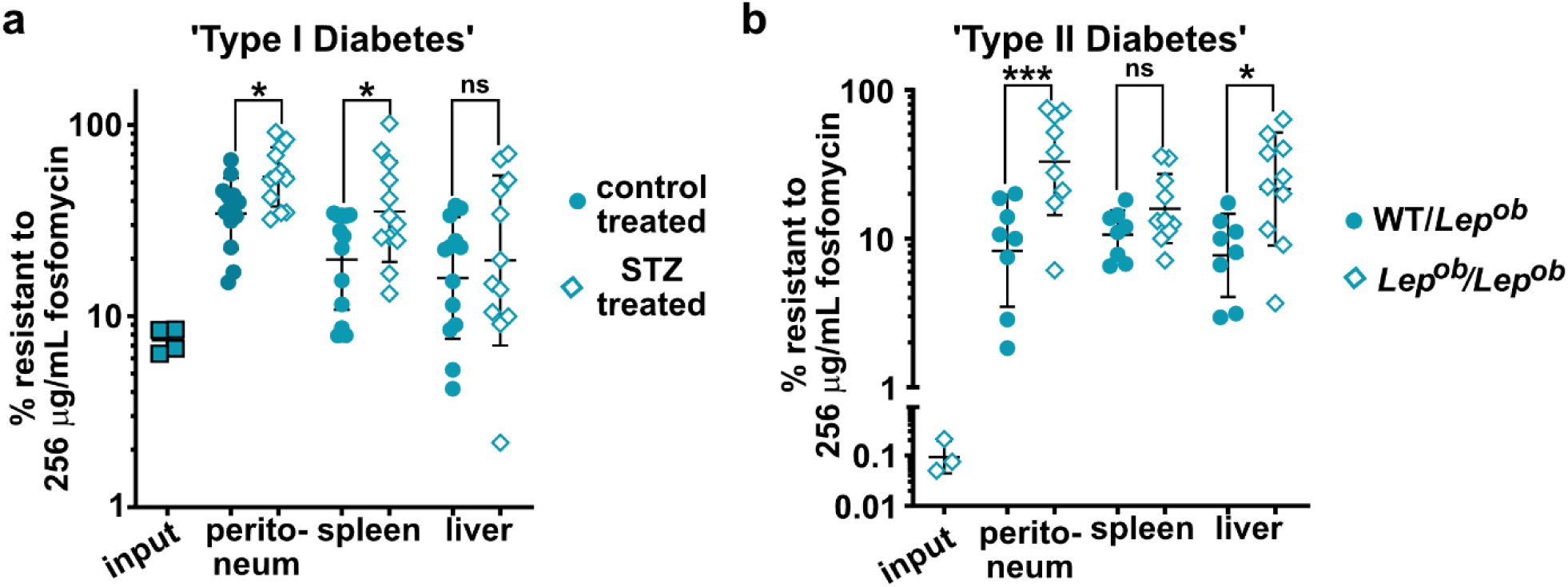
Glucose availability increases the frequency of the resistant subpopulation in murine models of diabetes/hyperglycemia. (a-b) Proportion of colonies resistant to fosfomycin in Mu208 plated directly from indicated tissues following infection of mice. Shown is geometric mean with geometric standard deviation. Mice were infected with ∼2x108 colony forming units for ∼24 h. (a) Prior to infection, mice were treated with control or streptozotocin (STZ), from three independent experiments with 13 mice total for control and 12 mice total for STZ treated. * indicates p= 0.0308 (peritoneum), 0.0390 (spleen) and ns indicates p=0.5076 by one-way ANOVA with Šídák’s multiple comparisons test, F (5, 69) = 6.762. (b) B6.Cg-*Lep*ob/J heterozygous or homozygous (*Lepob/Lepob*) mice were infected, from two independent experiments with 8 mice per total for WT/*Lepob* and 10 mice total for *Lepob/Lepob*. *** indicates p= 0.0002, ns indicates p= 0.6988, and * indicates p= 0.0229 by one-way ANOVA with Šídák’s multiple comparisons test, F (5, 47) = 6.756.

## DISCUSSION

We investigated the mechanism of fosfomycin heteroresistance in *Enterobacter cloacae* complex strain Mu208. While this isolate encodes the fosfomycin resistant determinant FosA, the majority of the cells are killed by the breakpoint concentration of fosfomycin. We found that the majority of the cells are susceptible at this concentration because they express sufficient GlpT, a fosfomycin importer. Why FosA is incapable of inactivating the intracellular fosfomycin at this concentration remains unclear. It appears that in wildtype cells grown in Mueller-Hinton broth, the balance of FosA activity and fosfomycin entry via GlpT results in homogenous resistance to low fosfomycin concentrations (64-128 µg/mL) but at breakpoint concentrations only a portion of cells are resistant, and only a portion of this population is resistant to 512 µg/mL fosfomycin. Our data suggests that *fosA* expression and/or synthesis of the FosA cofactor glutathione may be limiting, as expression from a multicopy plasmid of either *fosA* or *gshA* increases the frequency of the resistant population compared to wildtype (Figure 1g-h).

We found that the heterogeneity in fosfomycin resistance was derived from *glpT* promoter activity heterogeneity, rather than the product of gene amplification. The regulation of *glpT* is complex. Altering the carbon source environment affects *glpT* expression, resulting in shifts in fosfomycin entry and resistance. When glucose is present, it is imported by PtsG and converted to glucose-6-phosphate, reducing the phosphorylation of the PtsG partner protein Crr. This state of Crr results in dampened cAMP synthesis by CyaA, such that CRP is not active, and unable to activate *glpT* transcription. At the same time, GlpR represses *glpT,* resulting in poor fosfomycin entry into the cell and fosfomycin resistance (Figure 6a). In the absence of glucose, phosphorylated Crr would activate CyaA, activating CRP. CRP can activate *glpT* expression. If glycerol or glycerol-phosphate are present, GlpR will be additionally de-repressed, leading to high levels of *glpT* and fosfomycin import yielding low fosfomycin resistance (Figure 6b). These changes can occur globally and likely explain the overall shift in resistance profiles between, for instance, Mu208 grown in M9 glycerol versus M9 glucose (Figure 4e-f). On the other hand, for any one growth condition, heterogeneity of resistance exists in wildtype Mu208 (Figure 6c-d). We attribute this heterogeneity to a broad distribution in P*_glpT_* activity (Figure 2b), likely mediated by heterogeneity in CRP activity (Figure 3f). Together, heterogeneity in *glpT* expression results in population-wide differences in fosfomycin resistance.

**Figure 6.**
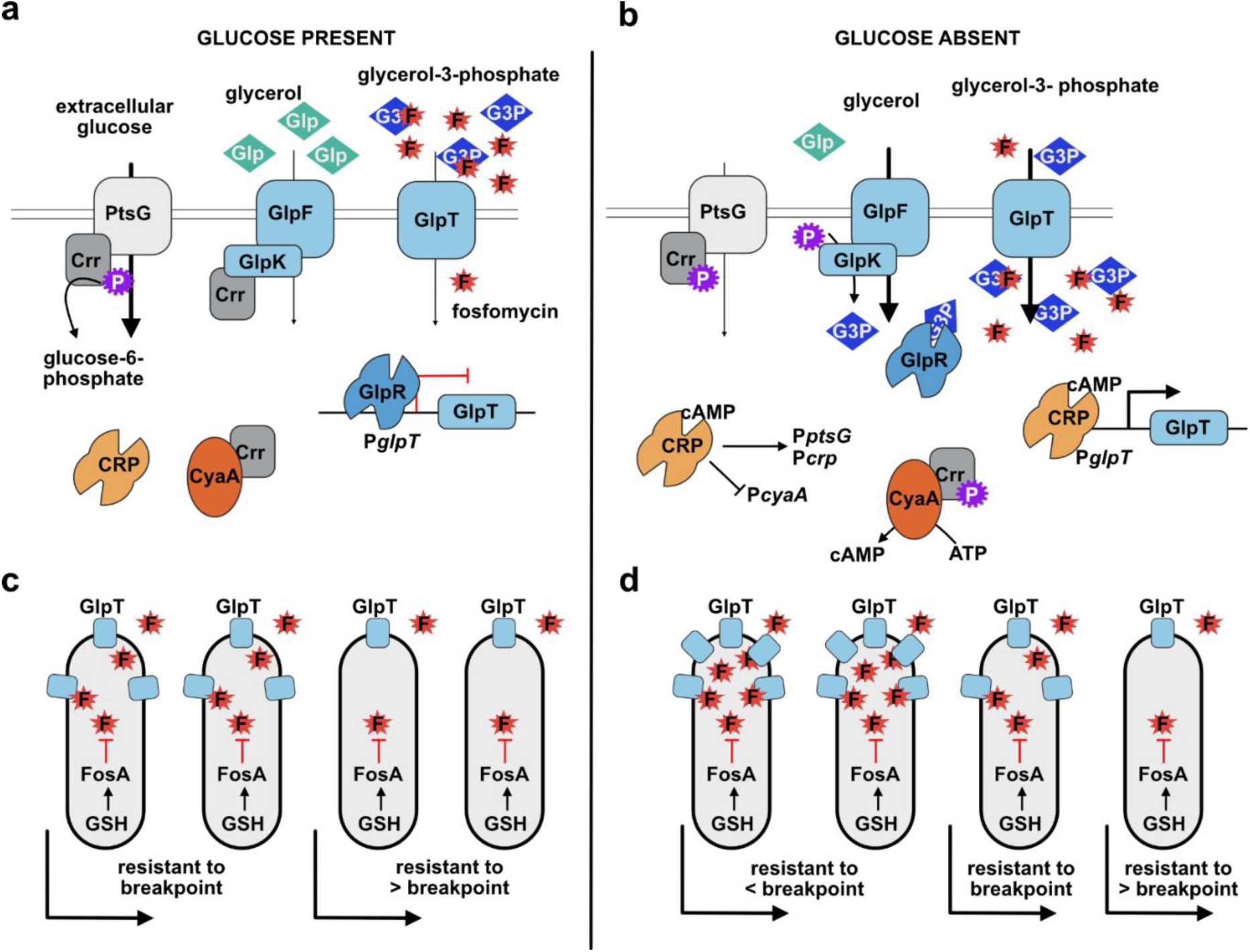
Proposed model of the effects of glucose availability on fosfomycin heteroresistance in *Enterobacter* Mu208. (a-b) Repression or activation of *glpT* transcription by the presence of glucose or glycerol carbon source is mediated through GlpR and CRP. (c-d) Population wide distribution of *glpT* expression generated resistant subpopulations. The effect of glucose shifted the entire population’s *glpT* expression low, but heterogeneity in *glpT* expression and relative fosfomycin resistance remained.

Fosfomycin HR presents challenges clinically. Depending on the frequency of the resistant subpopulation at the breakpoint concentration, isolates may be designated resistant or susceptible by conventional antimicrobial susceptibility testing. We showed that Mu208 can cause treatment failure in mice when treated with fosfomycin, and treatment of HR isolates can occur when the isolate is designated susceptible. Unfortunately, clinical fosfomycin susceptibility testing is fraught with confusion. Disk diffusion or Etest are often used. Resistant colonies are sometimes visualized within the zone of clearing on these fosfomycin susceptibility tests. However, there is disagreement about their relevance and how to interpret these colonies. In part because *glpT* mutants exhibit negative fitness costs ^17,31^, European (EUCAST) guidelines are to ignore these resistant colonies and classify such isolates as fosfomycin susceptible^32^. It is critical that the underlying biology of fosfomycin resistance be clearly understood to accurately guide effective therapy.

Our data additionally suggest that the host metabolic environment or co-morbidities could potentially affect fosfomycin therapy outcome. Because of the important role extracellular glucose played in repressing *glpT* expression and increasing population-wide fosfomycin resistance, we tested and found that excess host glucose in models of hyperglycemia/diabetes increased the frequency of the resistant subpopulation in strain Mu208. As the appreciation for the role of bacterial metabolism on antibiotic resistance grows^33^, our data highlight the potential for the host metabolic environment affecting bacterial metabolism and antibiotic resistance.

## MATERIALS AND METHODS

### Isolate information

*Enterobacter cloacae* complex strains Mu208, Mu819, Mu898, Mu909, Mu866, and Mu1197 were collected as part of the Georgia Emerging Infections Program as part of the CDC’s EIP Multi-site Gram-negative Surveillance Initiative (MuGSI) in Georgia, USA. *Enterobacter kobei* Mu208 is a clinical isolate previously described^15^. Strains and plasmids in this paper are detailed in Table 1.

**Table 1:**
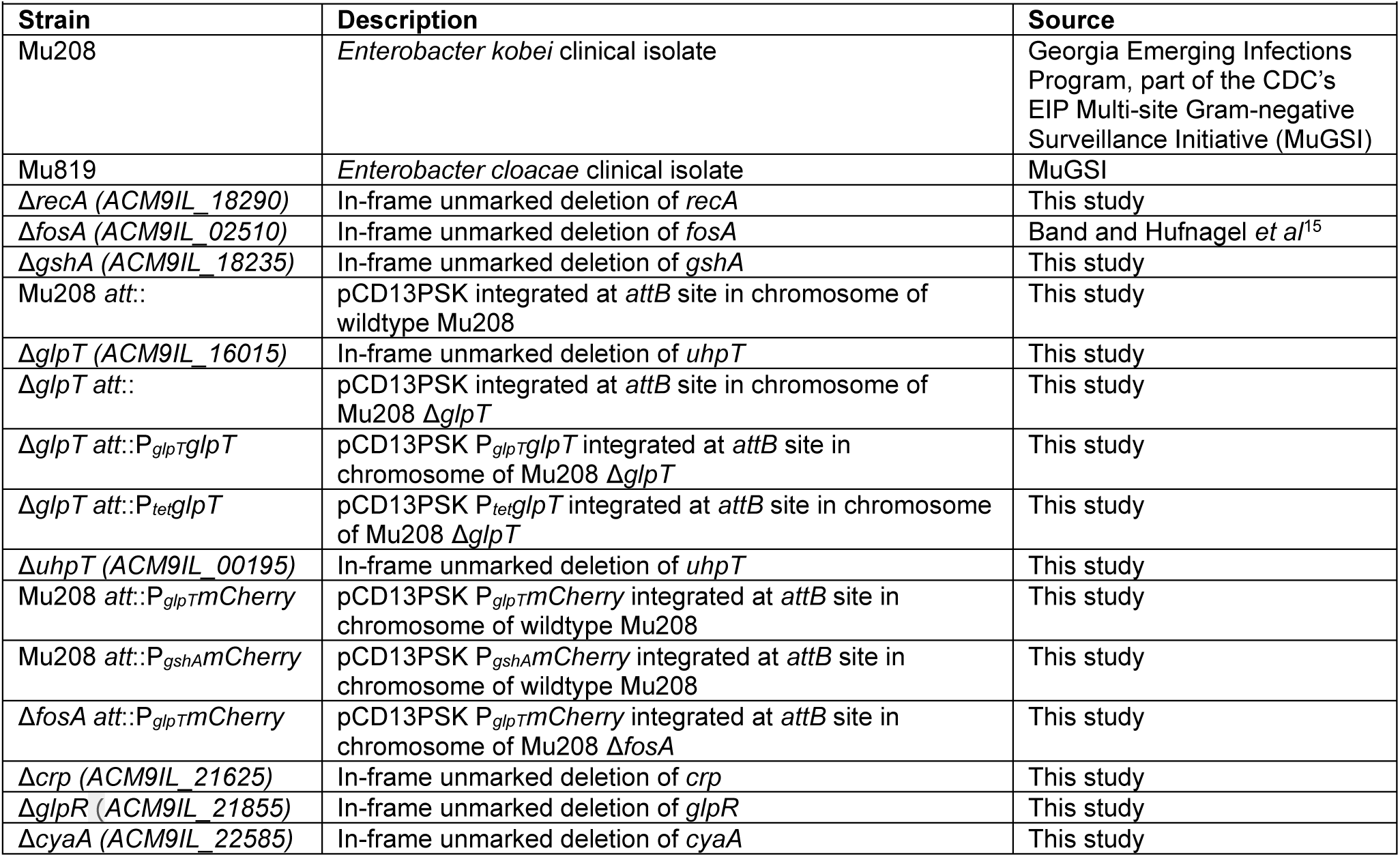
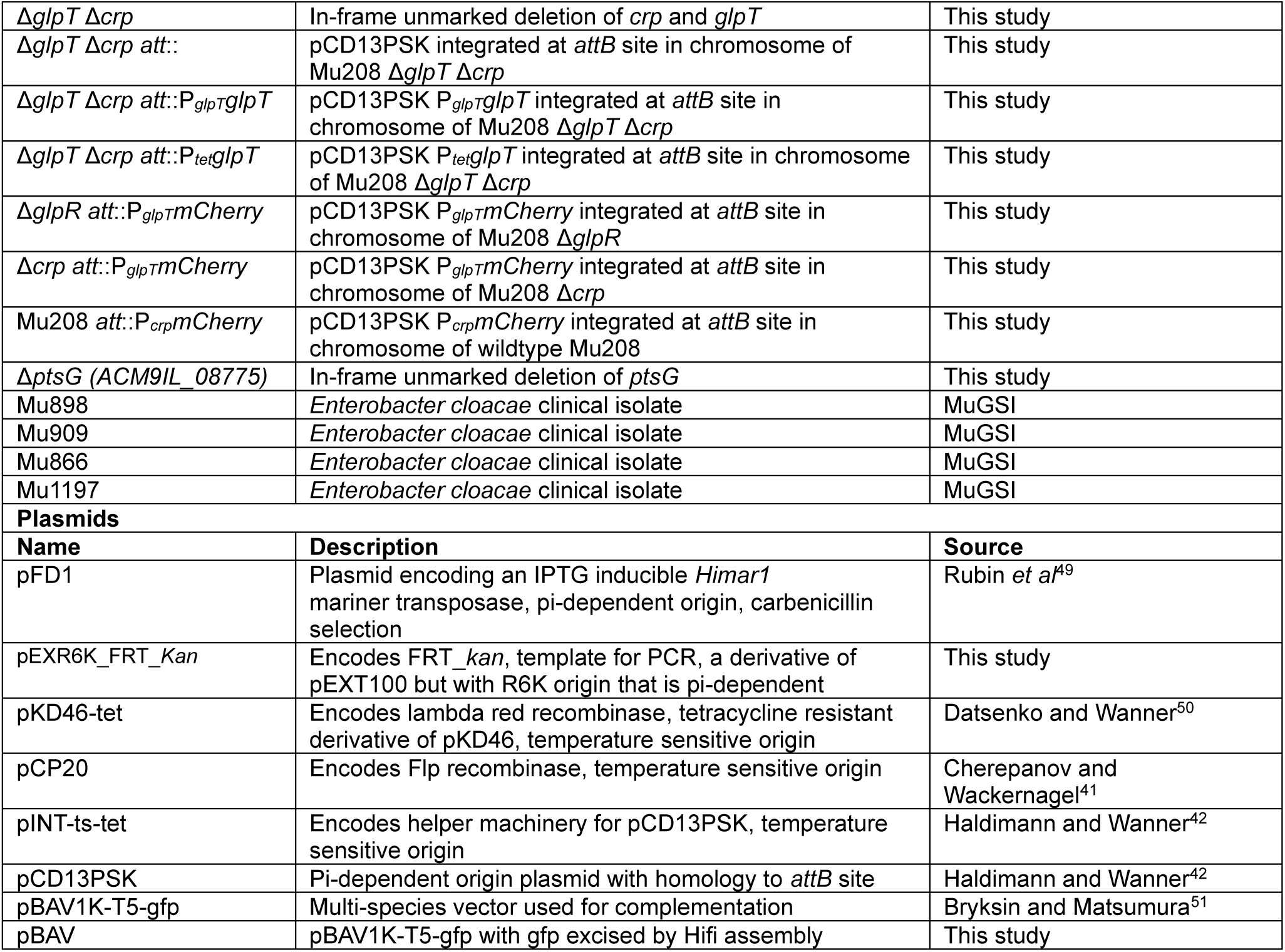
Strains and Plasmids.

### Reagents

Mueller-Hinton agar (MHA; BD Difco) and Mueller-Hinton broth (MHB; BD Difco) were used throughout as indicated. M9 broth was prepared with 1X M9 Salts (BD Difco), 100 µM CaCl_2_, 2 mM MgSO_4_ and carbon source as indicated, either glycerol (Promega #H5433) or anhydrous dextrose (VWR #BDH9230). For growth of *E. cloacae* before plating on MHA containing fosfomycin, bacteria were grown at 37°C with shaking, 1.5 mL volume in 12x75 mm polypropylene aeration tubes (Globe Scientific #110438). When MHB was used, 25 µg/mL glucose-6-phosphate was included per CLSI guidelines^34^. YESCA was made by autoclaving 1 g/L yeast extract (Fisher BP1422) and 10 g/L casamino acids (Acros #61204) in water. YESCA agar included 20 g/L agar (Sigma A1296).

Screening for GlpT function used M9 agar made by 2X M9 broth containing 2% glycerol phosphate disodium salt hydrate (Sigma # G6501) combined with 2X agar (Sigma A1296).

Antibiotics used were: fosfomycin disodium salt (TCI #F0889), kanamycin sulfate (Teknova #K2150; 35 µg/mL for strain Mu208), tetracycline (Alfa Aesar #J61714; 20 µg/mL for Mu208), chloramphenicol (Fisher #BP904, 50 µg/mL for Mu208) and spectinomycin dihydrochloride pentahydrate (Sigma #S4014; 50 µg/mL for *E. coli*, 125 µg/mL for Mu208).

Reduced L-Glutathione (Sigma #G4251), L-glutamic acid (Sigma #49449), and adenosine 3’,5’-cyclic monophosphate sodium salt monohydrate (Sigma A6885) were added at concentrations indicated.

Medium gel pads used for cell imaging under microscope: UltraPure^TM^ agarose (Invitrogen #16500100) and MHB was used to prepare MHB 1.5% agarose gel pad. M9 sole carbon source 10% polyacrylamide (PA) gel pad was made by mixing acrylamide (Bio-Rad #1610140, 10% (w/v)), N, N-methylene bisacrylamide (Bio-Rad #1610142, 0.15% (w/v)), ammonium persulfate (Bio-Rad #1610700, 0.75% (w/v)) and TEMED (Bio-Rad #1610800, 0.25% (w/v)), with M9+0.8% glycerol or M9+0.4% glucose. Polymerization of PA gel pad was performed under room temperature for 20 minutes^35^.

### Whole genome sequencing

Mu208 was plated on MHA containing 0 or 256 µg/mL fosfomycin Biomass was collected and subject to whole genome sequencing following extraction with Wizard Genomic DNA Purification kit (Promega). The genome of Mu208 grown without fosfomycin (mostly susceptible) was subjected to Illumina (650 Mbp) and Nanopore (300 Mbp ONT) sequencing to create a reference genome and quantify gene copy variation. The genome of the resistant population from growth on agar with fosfomycin was subject to Illumina sequencing and mapped to the respective reference genome with CNV analysis. Quality control and adapter trimming was performed with bcl2fastq^36^. Reads were mapped to their respective references via bwa mem^37^. PCR and optical duplicates were marked and excluded from the analysis using PicardTools’ ‘MarkDuplicates’^38^ functionality. Aligned read counts were imported into R’s CNOGpro^39^package. CNV events were called via CNOGpro using a bootstrapping method, which calculates an average gene number event giving a possible upper and lower bound. Genome sequencing and analysis were performed by SeqCenter (seqcenter.com).

### Population analysis profile

Population analysis profile (PAP) was performed as described previously^15^ and indicated in Supp. Fig.1. A given strain was grown overnight from a single colony streaked to MHA from -80°C glycerol stocks. After approximately 16-20 h of growth, the culture was diluted (1:250 for MHB, except 1:100 for strains with a growth defect [Δ*gshA*, Δ*crp*, Δ*cyaA*], and 1:100 for M9 broth) into fresh media and grown for 6 h in MHB or 7 h in M9. The cultures were serially diluted in PBS in a 96-well plate (Falcon) and 7.5 or 10 µl of each dilution was plated on MHA containing fosfomycin as indicated. Colonies were enumerated after 24-48 h of growth. For many experiments shown, the same protocol was followed but the cultures were plated only on MHA containing 0 or a single fosfomycin concentration indicated.

The surviving colonies are enumerated and the isolate is classified as resistant if at least 50% of the total colonies grow at 1 or 2X breakpoint. An isolate is considered susceptible if less than 0.0001% (-6 logs) of the cells grow at any concentration shown. An isolate is considered heteroresistant if there greater than 0.0001% survival at 1X and 2X breakpoint and an ≥8-fold difference between the resistance of the subpopulation and resistance of the main population. As an example for Mu208, if <50% of the population survives at 256 µg/mL, the resistance of the main population is 128 µg/mL. The resistance of the subpopulation that survives on 1024 µg/mL is greater than 1024 µg/mL, making the fold change ≥8. The limit of detection is approximately -7 logs but varies based on the density of the culture, in the figures the y-axis is set at -8 logs of survival and data points ≤-7 logs are generally at the limit of detection. See Supp. Fig. 1 for graphical representation.

In defining the features of heteroresistance in the isolates of this work, based on four guidelines set forth by Andersson et al^3^:

1. Clonality: these isolates demonstrate monoclonal heteroresistance, they are purified isolates and single colonies are used throughout experiments.
2. Level of resistance: the MIC of the resistant subpopulation in wildtype Mu208 is ≥8X the MIC of the main population, when comparing the amount of killing by the lowest concentration of fosfomycin used throughout, and the growth of the resistant subpopulation at 1024 µg/mL.
3. Frequency of the resistant subpopulation: We consider the frequency of the subpopulation at 2X the CLSI breakpoint, for the strains used throughout, the frequency is ∼0.01% for wildtype Mu208 following growth in MHB.
4. Stability: wildtype Mu208 demonstrate unstable heteroresistance. As shown in the passage experiments after selection, there is a significant reduction in the resistant population frequency within a single passage (∼10 generations per passage) in antibiotic free media.

### Time kill

Mu208 and Mu819 were streaked from -80 glycerol stocks to MHA for isolation and a single colony was used to start overnight cultures in MHB grown for 18 h. Subsequently, 1.5 µl of this overnight culture was added to 1.5 mL MHB containing 512 µg/mL fosfomycin. At the timepoints indicated, 100 µl of culture was removed, serially diluted in PBS and plated on MHA containing 0 or 512 µg/mL fosfomycin, grown for 24 h, and surviving colonies were enumerated.

### Resistance stability assays

Mu208 was streaked from -80°C glycerol stocks to MHA for isolation and a single colony was used to start overnight cultures in 1.5 mL of M9+0.8% glycerol phosphate and grown 15-20 h. 15 µl of the cultures were diluted into 1.5 mL of M9+0.8% glycerol phosphate of grown for 6 h. These cultures were diluted and plated MHA containing 0 or 256 µg/mL fosfomycin, the “baseline”. 1.5 µL of the cultures were added to fresh M9+0.8% glycerol phosphate containing 256 µg/mL fosfomycin and grown for 16 h. Following growth, the cultures were diluted and plated MHA containing 0 or 256 µg/mL fosfomycin, the “+fosfomycin”. 1.5 µL of the cultures were added to fresh M9+0.8% glycerol phosphate and grown for 20 h. Following growth, the cultures were diluted and plated MHA containing 0 or 256 µg/mL fosfomycin, the “drug-free media”.

### Murine treatment failure

∼8 week old female C57BL/6J mice (Jackson Laboratory) were injected intraperitoneally with 100 µl of Mu819, Mu208, or Mu208 Δ*gshA* suspended in sterile PBS at a density of ∼2x10^9^ CFU/mL. Following infection, mice were treated at 200 mg/kg using a sterile stock of 40 mg/mL fosfomycin disodium salt via intraperitoneal injection at 2, 8, 14, 20, 26, and 32 h post-infection. Mice were closely monitored during the 96 h experiment, with mouse weights being taken every 6 hours and mice were euthanized by carbon dioxide asphyxiation if moribund or below 80% starting weight, or at the conclusion of the experiment. All experiments were conducted in compliance with approved protocols and guidelines of the Emory University Institutional Animal Care and Use Committee (IACUC).

### Transposon screen

750 µl of a culture of a spontaneous streptomycin resistant variant of Mu208 was combined with 750 µl *E. coli* BW19851 carrying pFD1, collected by centrifugation and resuspended in 50 µl YESCA. The suspension was plate on a 0.45 µM nitrocellulose membrane placed on top of a YESCA agar plate and incubated at 37°C for 2.5 h. The nitrocellulose was washed in a 15 ml tube with YESCA containing 1 mM IPTG and 100 µg/mL streptomycin, grown for 3 h at 37°C, diluted and plated on MHA containing 100 µg/mL streptomycin and 35 µg/mL kanamycin. After 24 h of growth at 37°C, colonies were replica-plated using sterile velvet onto MHA containing kanamycin or kanamycin and 256 µg/mL fosfomycin. Colonies that appeared on kanamycin but not kanamycin+fosfomycin were restreaked for isolation. To identify the transposon insertion site, colony PCR was performed using primers 71/72 to yield the DNA flanking the tranpsosase. 2 µl of the PCR product from JCP71/72 was the template for subsequent nested PCR with JCP73/74. PCR products were Sanger sequenced (Azenta) using JCP74 and aligned to the Mu208 genome.

### Strain construction: gene deletions

Lambda-red based allelic exchange was used to replace the coding sequence of genes with a kanamycin resistance gene, and then the gene was removed with Flp recombinase^40,41^ to create unmarked in-frame deletions. The kanamycin resistance gene from pEXR6K_kanFRT was cloned using Promega GoTaq 2X master mix with Flp recognition sequence and homology to regions flanking the gene to be replaced using primers DHP175/181 (*recA*), DHP65/66 (*fosA*), DHP109/110 (*gshA*), DHP3/4 (*glpT*), DHP5/6 (*uhpT*), DHP49/50 (*crp*), JCP495/496 (*glpR*), DHP51/52 (*cyaA*), and DHP127/128 (*ptsG*), detailed in Supp. Table 3. The purified PCR product was electroporated into competent Mu208 (or mutant) carrying pKD46-tet and transformants were selected on kanamycin. Transformants were re-streaked to kanamycin for isolation and subject to PCR for successful allelic exchange as indicated by a product size change using primers flanking the gene to be replaced: primers DHP179/182 (*recA*), DHP73/74 (*fosA*), DHP117/118 (*gshA*), DHP9/10 (*glpT*), DHP11/12 (*uhpT*), DHP57/58 (*crp*), JCP497/498 (*glpR*), DHP59/60 (*cyaA*), and DHP135/136 (*ptsG*), detailed in Supp. Table 3. The mutants were then made electrocompetent and electroporated with pCP20^41^. Transformants were selected at 30°C on chloramphenicol, patched to chloramphenicol at 30°C and grown for 24 h, then patched to MHA and MHA+kanamycin. Kanamycin-sensitive mutants were streaked for isolation and PCR was used with the same flanking primers to screen for the loss of the kanamycin resistance gene.

### Strain construction: plasmid-borne complementation

The coding sequences of the following Mu208 genes were cloned into pBAV-T5 at BamHI and NcoI sites using primers DHP290/291 (*fosA*), DHP286/287 (*gshA*), DHP284/285 (*crp*). Using Hi-fi assembly (NEB), pBAV-T5 amplified by PCR using JCP280/281 was assembled with *glpR* (JCP499/500), cyaA (TO131/132), and *ptsG* (TO142/143). Plasmids were propagated in *E. coli* NEB5α, Sanger sequenced (Azenta) using primers JCP339/340, and electroporated into appropriate strain with kanamycin selection.

### Strain construction: chromosomally integrated plasmids

pCD13PSK was digested with BamHI and SacI. *P_glpT_glpT* was amplified from the chromosome with primers (JCP477/478) that included homology to digested pCD13SPK and combined via NEB Hifi assembly. Similarly, *glpT* and its 5’ UTR were amplified with primers (JCP476/477) which included homology to pCD13PSK, and the forward primer (JCP476) included P*_tet_* to generate pCD13PSK *P_tet_glpT.* JCP511/512 amplified a linear product of pCD13PSK *mCherry* (Emily Crispell, Weiss laboratory) for Hifi assembly with Mu208 promoters to generate pCD13PSK P*_gene_mCherry:* primers amplified the promoter with homology to pCD13PSK *mCherry*: P*_glpT_* (JCP513/514), P*_gshA_* (JCP515/516), and P*_crp_* (JCP521/522).

Plasmids were propagated in *E. coli* PIR2 with spectinomycin, and Sanger sequenced (Azenta) using primers JCP49/585. Mu208 wildtype or indicated mutants were made electrocompetent, transformed with pINT-ts-tet and selected for with tetracycline. Strains carrying pINT-ts-tet were made electrocompetent and transformed with appropriate pCD13PSK plasmid. Of note, following 1 h of recovery at 37°C, the culture was moved to 42°C for 30 min before plating on spectinomycin.

Integration of pCD13PSK at the *attB* site was confirmed by colony PCR using JCP474/53. To additionally confirm, the region of the chromosome encoding the cloned genes and pCD13PSK with amplified with JCP474/539 and Sanger sequenced (Azenta) using primers JCP49/585. For experiments, spectinomycin was not used because the integration is stable^42^.

### Imaging of cells and mCherry expression

Mu208 wildtype carrying *att*:: P*_glpT_mCherry*, *att*:: P*_gshA_mCherry*, and *att*:: P*_crp_mCherry*, along with the indicated Mu208 mutants carrying P*_glpT_mCherry* were used to visualize population-level variation in promoter activation prior to fosfomycin exposure, and to assess viability following treatment.

Imaging of cells and mCherry expression was performed as previously described^43,44^. Once a cell culture reached OD_600_ ∼ 0.1, cells were placed on a 35 mm glass-bottom Petri dish (Cellvis) and covered with an MHB 1.5 % agarose gel pad, or 10 % Polyacrylamide gel pad containing M9+glucerol or M9+glucose, as needed. Cells were imaged using an inverted microscope (Olympus IX83) with a 60× oil immersion phase-contrast objective lens, housed within a pre-warmed (37 °C) incubator chamber (InVivo Scientific). mCherry fluorescence was detected using a TRITC filter cube.

Snapshot imaging of cells was performed before exposure to fosfomycin. Time-lapse images of cells were captured every 30 minutes after adding fosfomycin to the agarose pad covering cells.

### Image analysis

Fluorescence intensity of mCherry was measured using a plug-in for Fiji ImageJ software, named MicrobeJ^45^ (version 5.13 I (22)). Reported intensities are the results after subtracting extracellular background fluorescence as well as autofluorescence of wildtype Mu208 cells from intracellular fluorescence signals.

Viability of cells were tracked from time-lapse images after fosfomycin exposure. Continuous cellular division and colony formation represented survival. Killing of cells by fosfomycin was defined as visible lysis or permanent arrest of cell growth.

### qPCR

For cAMP addition, Mu208 was grown overnight in M9+0.8% glycerol and 15 µl was added to 1.5 mL fresh M9+0.8% glycerol with and without 5 mM cAMP. After 7 h of growth, 1.3 mL was added to 2.6 mL RNAprotect (Qiagen #76104). RNA was subsequently isolated using RNeasy Mini Kit (Qiagen #74104) then treated with DNAse (BioLab #M0303S) according to manufacturer’s instructions. Reverse-transcriptase quantitative PCR was performed with Power SYBR Green RNA to C_T_ 1-Step Kit (Applied Biosystems #4389986). *rplB* was detected with TO168/169 and was used as the housekeeping for normalization, *glpT* with DHP264/265. The fold change for each biological replicate was compared to the same replicate from no cAMP addition, resulting in a value of 1 for each no cAMP replicate. Fold change was calculated using the 2^-ΔΔCT^ method^46^.

For glucose addition, Mu208 was grown overnight in MHB and 2.5 µl was added to 2.5 mL MHB with or without 2% glucose added, and grown until OD reached ∼0.4-0.5. 1 mL was added to 2 mL RNAprotect, RNA was isolated as above, and qPCR performed as above.

### GlpT function phenotypic screening

Colonies from the strains and agar containing fosfomycin concentrations indicated in Supplementary Figure 6d were patched to M9 agar containing 2% glycerol phosphate. Mu208 wildtype and Δ*glpT* were patched as quality controls, wildtype grew but Δ*glpT* was unable to grow. Mu208 forms small and large resistant colonies on fosfomycin, and from our screening we identified that only the large colonies exhibited the GlpT-null phenotype. By patching the more frequent small resistant colonies and less frequent large resistant colonies, we calculated the frequency of the GlpT null phenotype as reported in Supplementary Figure 6d.

### Murine diabetic models

Streptozocin treated and control treated male C57BL/6J mice were sourced from Jackson Laboratory, Stock # 033853. Jackson Laboratory verifies a non-fasted blood glucose exceeding 250 mg/dL prior to shipment, but most mice have glucose exceeding 350 mg/dL^47^. These mice were infected intraperitoneally at ∼10 weeks of age with 100 µl of Mu208 suspended in sterile PBS at a density of 1.7-2.4x10^9^ CFU/mL.

B6.Cg-*Lep^ob^*/J heterozygous and homozygous male mice were sourced from Jackson Laboratory (Strain #000632). At 8 weeks male *Lep^ob^*/*Lep^ob^*mice have a mean non-fasted blood glucose of ∼350 mg/dL and WT/*Lep^ob^*mice have a mean of ∼200 mg/dL^48^. These mice were infected intraperitoneally at ∼8 weeks of age with 100 µl of Mu208 suspended in sterile PBS at a density of ∼2x10^9^ CFU/mL.

For both series of experiments, after 24 h, mice were euthanized by carbon dioxide asphyxiation. The peritoneal cavity was washed with 3 mL sterile PBS using a syringe and 26 gauge needle, and the liver and spleen were weighed and collected in 1 mL PBS. Organs were homogenized using a BioSpec Tissue Tearor (#98570-395). The suspensions were diluted in PBS, spread, and spot-plated on MHA containing 0 or 256 µg/mL fosfomycin for enumeration. All experiments were conducted in compliance with approved protocols and guidelines of the Emory University Institutional Animal Care and Use Committee (IACUC).

### Statistics

The number of experiments and the number of biological replicates are detailed in each figure legend, along with statistical test as appropriate. Unless otherwise specified, the mean is shown and error bars indicate standard deviation.

## MATERIALS AND DATA AVAILABILITY

Materials and underlying data are available upon request to David S. Weiss. Mu208 genome sequences are available at NCBI Genbank with accession numbers CP182179, CP182180, and CP182181.

## ACKNOWLEDGEMENTS

We would like to thank the US Centers for Disease Control and Prevention (CDC)’s Emerging Infections Program and the Georgia Multi-Site Gram-negative Surveillance Initiative (MuGSI), including Jesse Jacob and Monica Farley, for providing isolates. pBAV1K-T5-gfp was a gift from Ichiro Matsumura (Addgene plasmid # 26702 ; http://n2t.net/addgene:26702 ; RRID:Addgene_26702). We acknowledge the contribution of genome sequencing by SeqCenter. This work was supported by the National Institutes of Health [AI158080, AI141883, AI32956] and the National Institutes of Health’s Office of the Director, Office of Research Infrastructure Programs, P51 OD011132. DSW is supported by a Burroughs Wellcome Fund Investigators in the Pathogenesis of Infectious Disease award. The Multi-Site Gram-negative Surveillance Initiative is funded by the CDC. The content is solely the responsibility of the authors and does not necessarily reflect the official views of the National Institutes of Health

## SUPPLEMENTAL

**Supplemental Figure 1.**
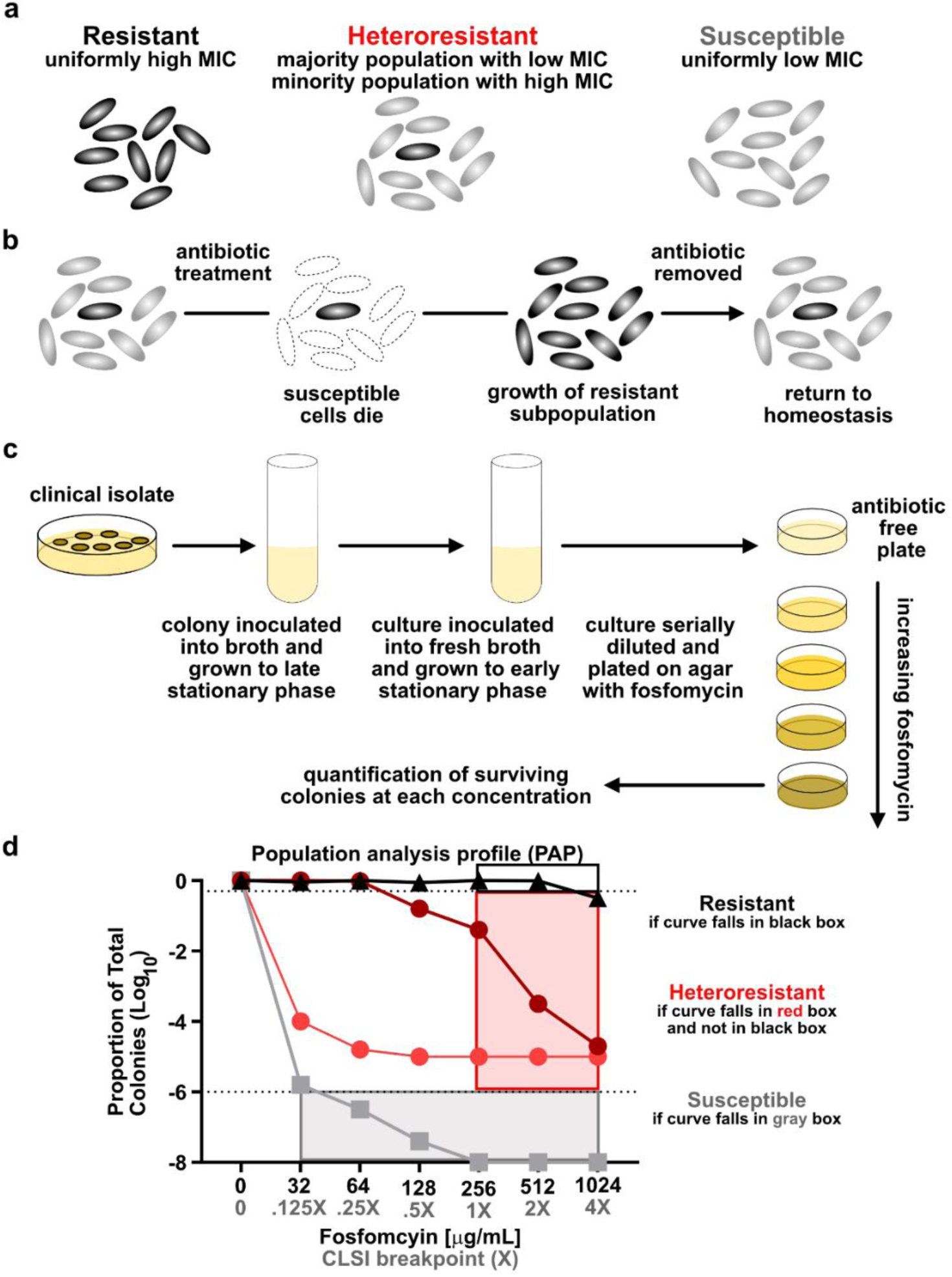
Overview of heteroresistance and population analysis profile (PAP). (a) A depiction of the cells grown from a single colony of an isolate exhibiting conventional resistance, heteroresistance, or susceptibility to a given antibiotic. (b) Population dynamics of a heteroresistant isolate following antibiotic treatment. (c) Fosfomycin PAP: a clinical isolate is isolated and grown in broth, then subcultured into fresh broth and grown to early stationary phase. The culture is then serially diluted, plated on agar with increasing concentrations of fosfomycin, and incubated. (d) The surviving colonies are enumerated and the proportion of total colonies that survive on a given concentration is calculated 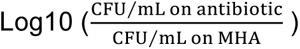. The isolate is classified as resistant if at least 50% of the total colonies grow at 1 or 2X breakpoint. An isolate is considered susceptible if less than 0.0001% (-6 logs) of the cells grow at any concentration shown. An isolate is considered heteroresistant if there is greater than 0.0001% survival at 1X and 2X breakpoint and an ≥8-fold difference between the resistance of the subpopulation and resistance of the main population. As an example, if <50% of the population survives at 128 µg/mL, the resistance of the main population is 64 µg/mL. The resistance of the subpopulation that survives on 512 µg/mL is greater than 512 µg/mL, making the fold change ≥8. Along the x-axis, the breakpoints based on CLSI are shown in gray, and the corresponding concentration of fosfomycin is in black.

**Supplemental Figure 2.**
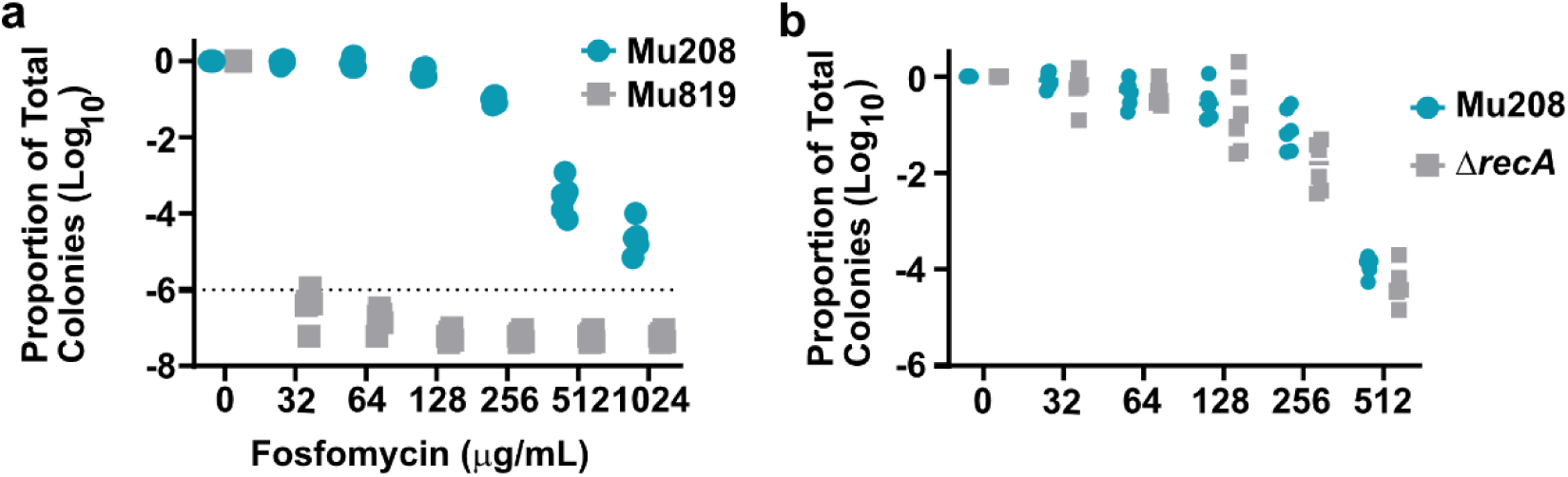
Fosfomycin heteroresistance in Mu208 is *recA*-independent. (a) Alternative presentation of population analysis profile (PAP) of Mu208 and Mu819, from three independent experiments with 2 biological replicates each. (b) Population analysis profile (PAP) of Mu208 and the Δ*recA* deletion strain, from two independent experiments with 3 biological replicates each.

**Supplemental Figure 3.**
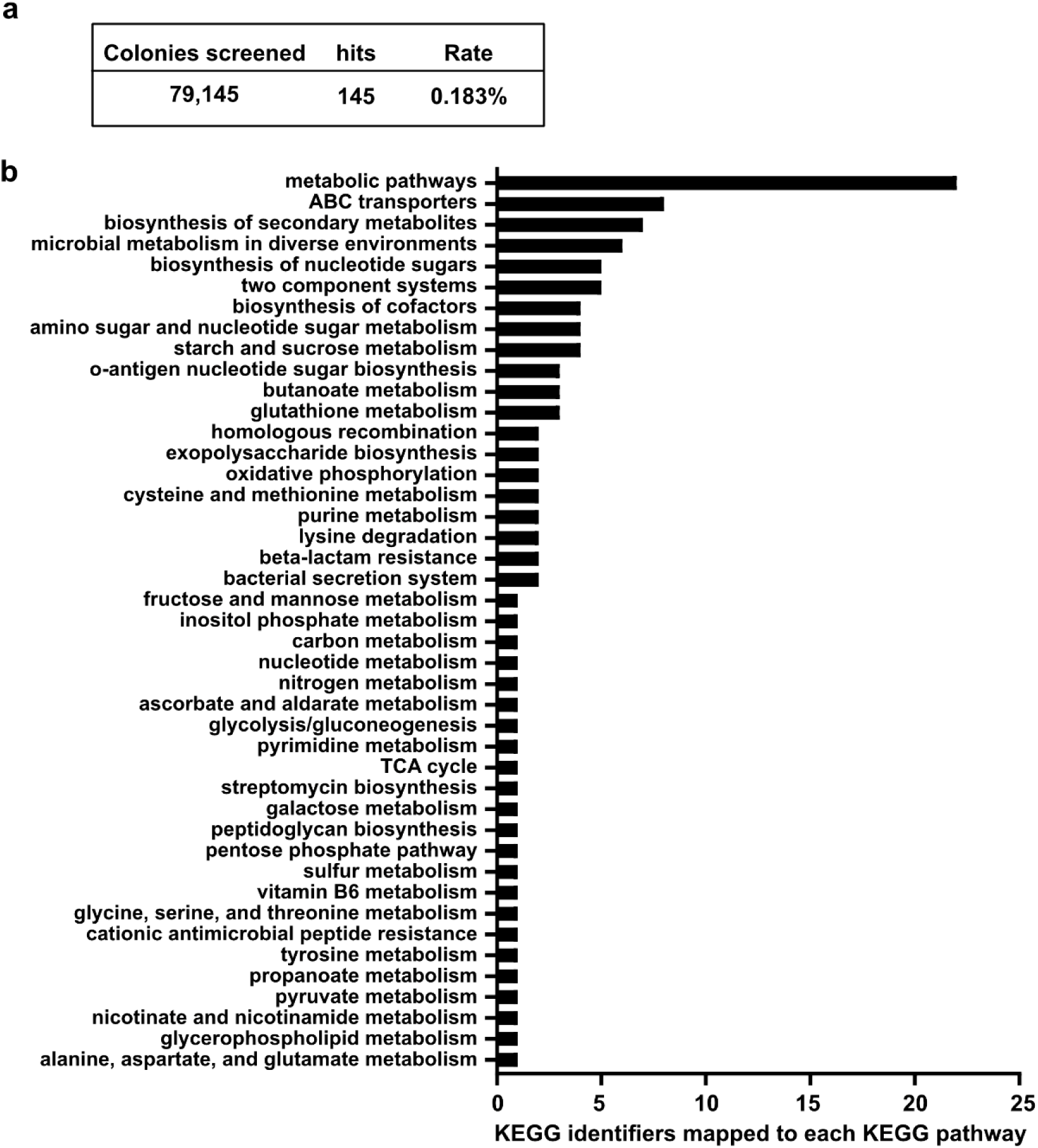
Transposon screen for loss of fosfomycin resistant subpopulation identifies key role for metabolic genes. (a) Results of transposon screen. (b) The KEGG identifiers of each transposon-disrupted gene in the screen was mapped to KEGG pathways as above. A portion of the genes did not map to a KEGG identifier/pathway. Full list is provided in Supplemental Table 2

**Supplemental Figure 4.**
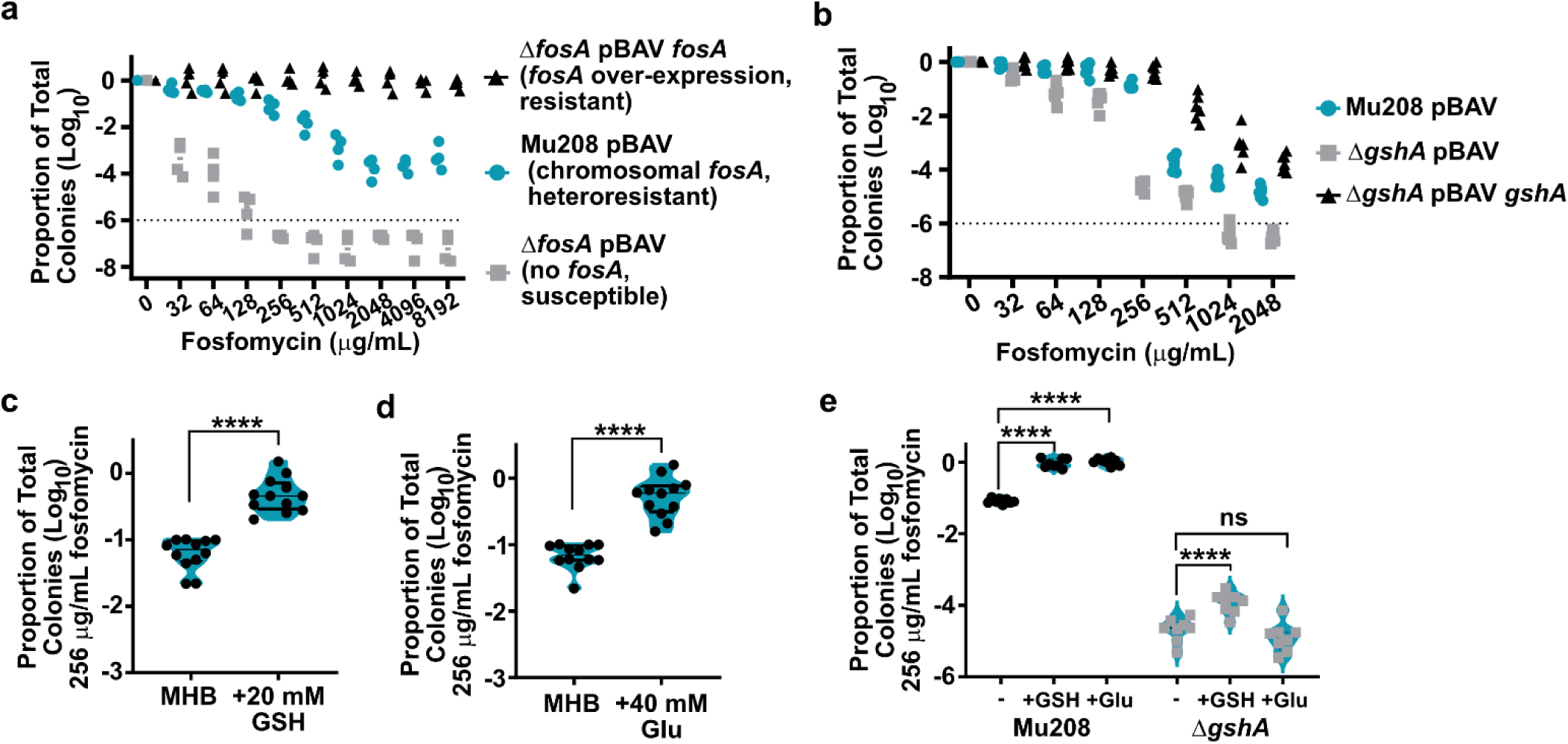
FosA and glutathione synthesis support the resistant subpopulation. (a) Alternative presentation of PAP of Mu208, the Δ*fosA* deletion strain, and its complement two independent experiments with 2 biological replicates each. (b) Alternative presentation of PAP of strains Mu208, Δ*gshA* mutant and its complement two independent experiments with 3 biological replicates each. (c) Proportion of strain Mu208 population resistant to fosfomycin following growth in Mueller Hinton broth, with addition of 20 mM glutathione (+GSH), from three independent experiments with 4 biological replicates each. **** indicates p<0.0001, t=8.946, df=11 (d) Proportion of population resistant to fosfomycin following growth in Mueller Hinton broth, with addition of 40 mM L-glutamic acid (+Glu) in strain Mu208 from three independent experiments with 4 biological replicates each. Some of the MHB replicates are the same between c and d. **** indicates p<0.0001 by paired t-test, t=7.568, df=11. (e) Proportion of population resistant to fosfomycin following growth in Mueller Hinton broth, with addition of 20 mM glutathione (+GSH) or 40 mM L-glutamic acid (+Glu) in strains Mu208 and Δ*gshA*, from two independent experiments with 4 biological replicates each. **** indicates p<0.0001 by one-way ANOVA with Šídák’s multiple comparisons test, F (5, 42) = 669.8.

**Supplemental Figure 5.**
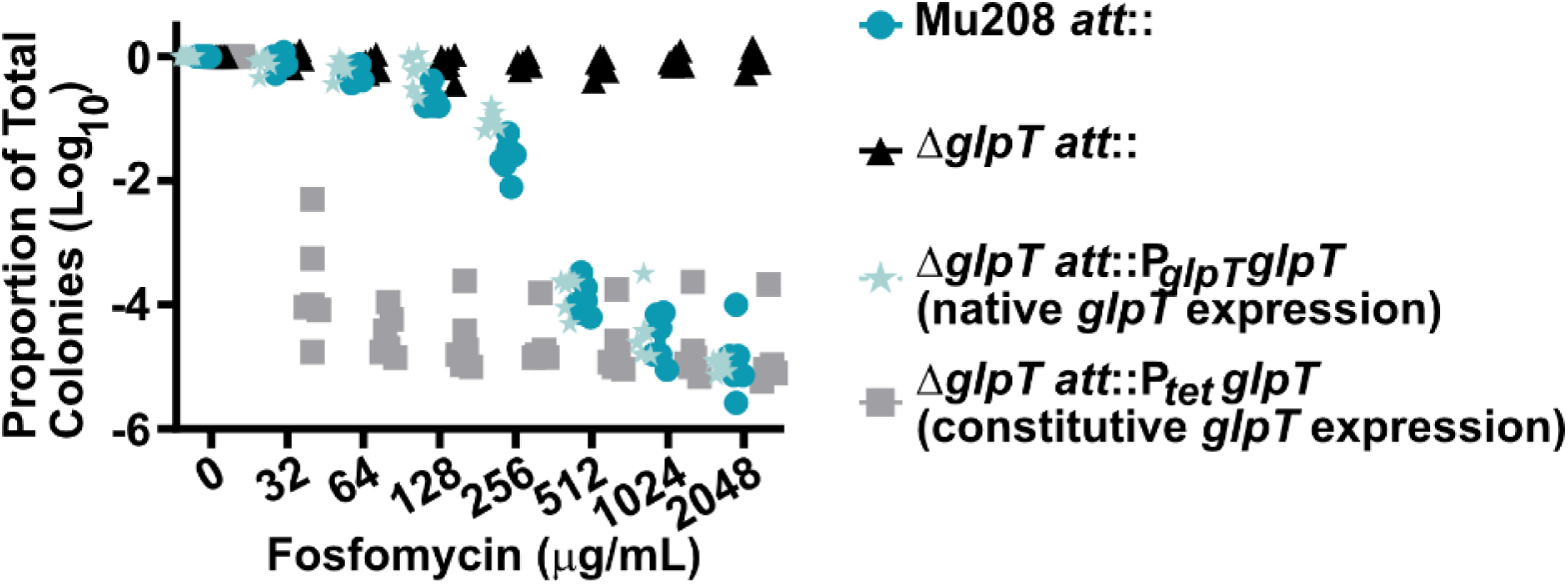
GlpT expression controls subpopulation frequency. Alternative PAP presentation of fosfomycin heteroresistance in Figure 2a, from two independent experiments with 3 biological replicates each.

**Supplemental Figure 6.**
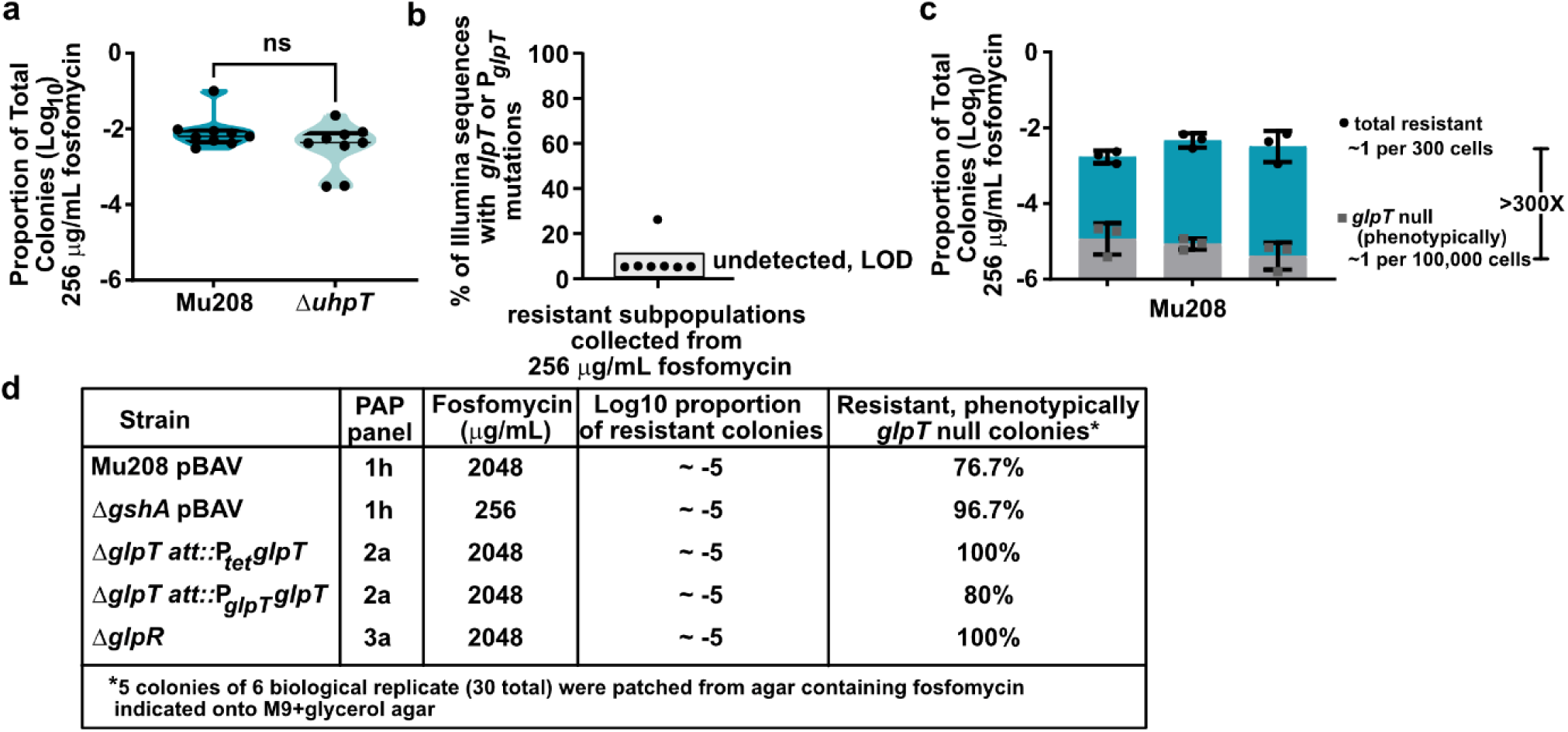
Spontaneous *glpT* mutants constitute a small portion of the resistant subpopulation. (a) Proportion of colonies resistant to fosfomycin in strains Mu208 and Δ*uhpT*, encoding the hexose-phosphate transporter, from three independent experiments with 3 biological replicates each. ns indicates p>0.05 by two-tailed t-test t=1.301, df=8. (b) P*glpTglpT* mutations detected in fosfomycin resistant colonies by variant calling from Illumina sequencing. 7 colonies collected from agar containing 256 µg/mL fosfomycin were sequenced. (c) Proportion of colonies resistant to fosfomycin and then screened for loss of GlpT function in Mu208, from three independent experiments (each bar) with 3 biological replicates total. (d) Fosfomycin resistant colonies from strains indicated where screened for GlpT function by assessing growth on M9+glycerol agar.

**Supplemental Figure 7.**
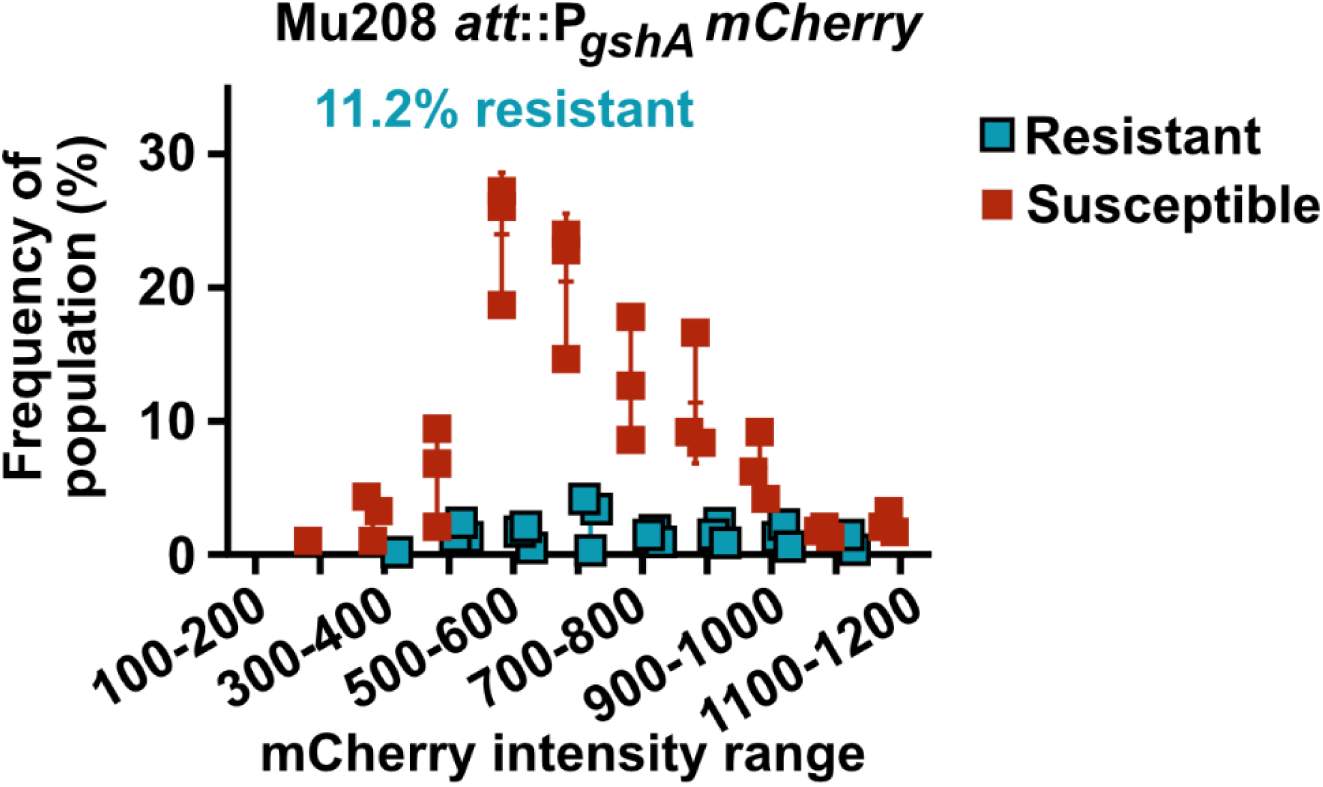
P*_gshA_mcherry* is not a predictor of survival. P*gshAmcherry* expression intensity among cells that survived or were killed by 128 µg/mL fosfomycin, from three independent experiments; data were determined from 700-1000 cells for each experiment.

**Supplemental Figure 8.**
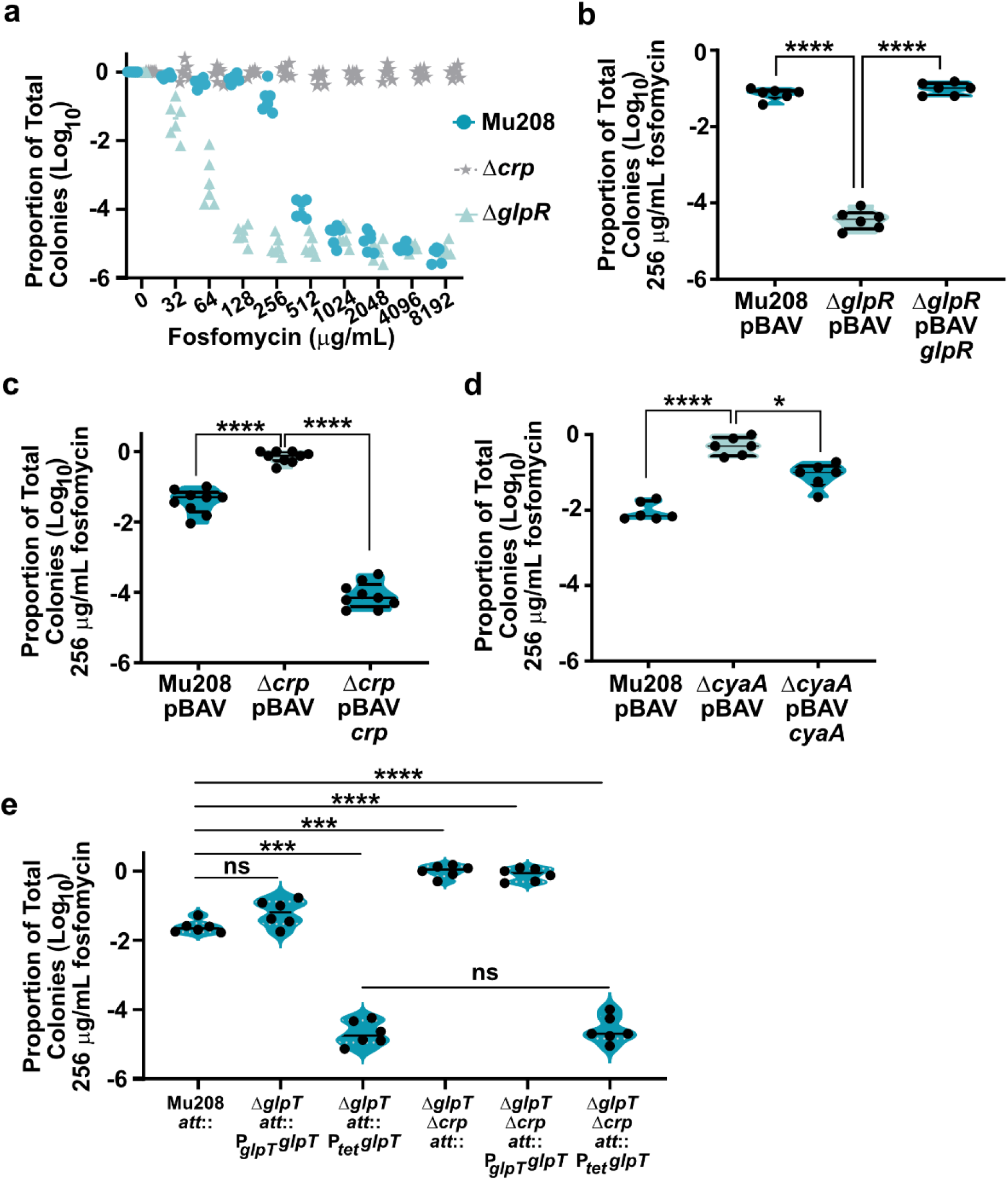
Regulation of *glpT* by GlpR and CyaA/CRP controls the subpopulation. (a) Alternative presentation of PAP for strains Mu208, Δ*cyaA*, Δ*crp*, and Δ*glpR*, from Figure 3a, from two independent experiments with 3 biological replicates each. (b-d) Proportion of colonies resistant to fosfomycin in strains Mu208, mutants, and their complement, from two independent experiments with 3 biological replicates each. * indicates p= 0.0127 and **** indicates p<0.0001 by RM one-way ANOVA with Dunnett’s multiple comparisons test, b: F (1.760, 8.799) = 799.0, c: F (1.208, 9.665) = 293.3, and d: F (1.411, 7.055) = 55.80. (e) Proportion of colonies resistant to fosfomycin in strains indicated, from two independent experiments with 3 biological replicates each. ns indicates p>0.05, *** indicates p= 0.0003 or 0.0001 (left to right) and **** indicates p<0.0001 by RM one-way ANOVA with Dunnett’s multiple comparisons test, F (2.748, 13.74) = 319.7.

**Supplemental Figure 9.**
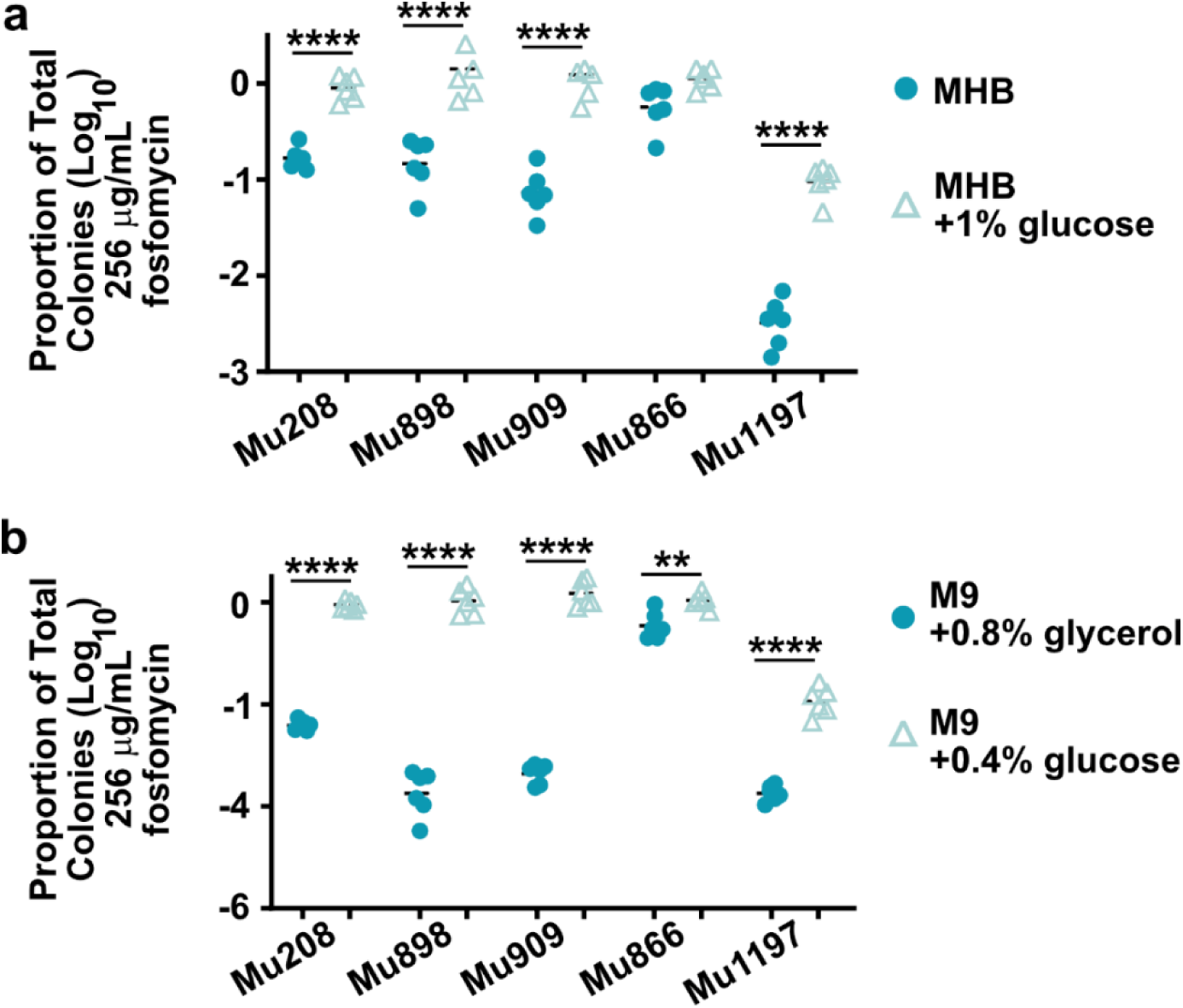
Glucose enhances the resistant subpopulation in additional carbapenem-resistant Enterobacter clinical isolates. Proportion of colonies resistant to fosfomycin following growth in (a) Mueller Hinton broth ±1% glucose and (b) M9 media with glycerol or glucose as the sole carbon sources. (a-b) from two independent experiments with 3 biological replicates each. ** indicates p<0.01 and **** indicates p<0.0001 by one-way ANOVA with Sidak’s correction for multiple comparisons, F (9, 50) = 87.55 for a and F (9, 50) = 302.2 for b.

**Supplemental Figure 10.**
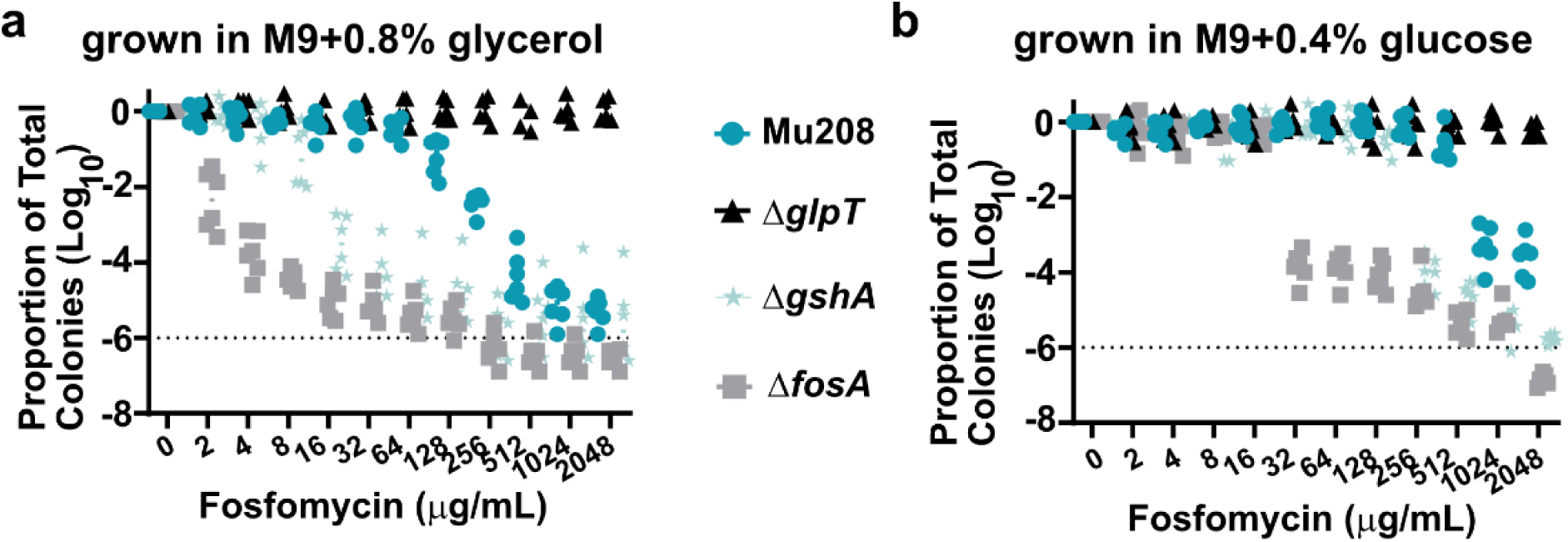
Glucose enhances the resistant subpopulation in Mu208. Alternative presentation of population analysis profiles from strains Mu208, Δ*glpT*, Δ*gshA*, and Δ*fosA*. (a) following growth in M9+ glycerol as the sole carbon source or (b) following growth in M9+glucose as the sole carbon source, from two independent experiments with 3 biological replicates each.

**Supplemental Figure 11.**
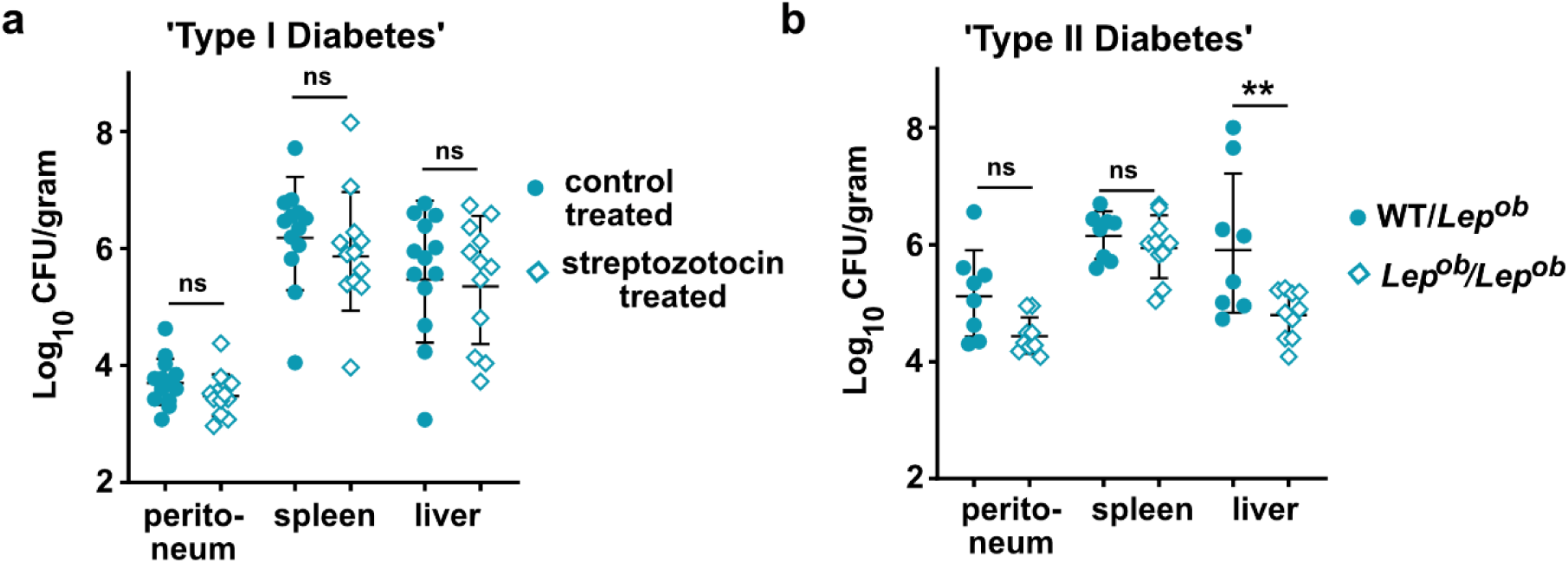
Total bacterial burdens in murine models of diabetes. Total colony forming units isolates from indicated tissue following infection with Mu208. Shown is geometric mean with geometric standard deviation Mice were infected with 1.5-2x10^8^ colony forming units for ∼24 h. (a) Prior to infection, mice were treated with control or streptozotocin (STZ), from three independent experiments with 13 mice total for control and 12 mice total for STZ treated. ns indicates p>0.05 by one-way ANOVA with Šídák’s multiple comparisons test, F (5, 69) = 24.14. (b) B6.Cg-*Lep*^ob^/J heterozygous or homozygous (*Lep^ob^/Lep^ob^*) mice were infected, from two independent experiments with 8 mice per total for WT/*Lep^ob^* and 10 mice total for *Lep^ob^/Lep^ob^*. ns indicates p>0.05, and ** indicates p=0.0013 by one-way ANOVA with Šídák’s multiple comparisons test, F (5, 47) = 10.30.

